# Active fluctuations modulate gene expression in mouse oocytes

**DOI:** 10.1101/347310

**Authors:** Maria Almonacid, Stephany El-Hayek, Alice Othmani, Isabelle Queguiner, Fanny Coulpier, Sophie Lemoine, Leila Bastianelli, Christophe Klein, Tristan Piolot, Philippe Mailly, Raphaël Voituriez, Auguste Genovesio, Marie-Hélène Verlhac

**Author notes:** correspondence should be addressed to AG and MHV.

## Abstract

In mammals, the nucleus is central in oocytes, not defining the future embryo axis. Nucleus centring depends on an F-actin mediated pressure gradient. In *Fmn2*^*−/−*^ oocytes, lacking the F-actin nucleator Formin 2, the nucleus is off-centre and can be centred by re-expressing Formin 2. Here, we addressed the biological significance of nucleus positioning in mammalian oocytes. Using a dedicated computational 3D imaging approach, we observed nuclear architecture alterations in mouse *Fmn2*^*−/−*^ oocytes. RNA sequencing of control versus *Fmn2*^*−/−*^ oocytes detected 2285 mis-regulated genes. Rescue experiments showed that the process of nuclear positioning impacts nuclear architecture and gene expression. Using signal processing methods coupled to biophysical modelling allowing the extraction of *in vivo* mechanical properties of the nuclear envelope, we showed that F-actin-mediated activity promotes nuclear envelope shape fluctuations and chromatin motion. We thus propose a mechano-transduction model whereby nucleus positioning via microfilaments modulates oocyte transcriptome, essential for further embryo development.

## Introduction

The position of the nucleus inside the cell can exert a morphogenetic influence conveying spatial and temporal information. A failure to correctly position the nucleus can promote disease, such as when nuclei from cortical neurons in the brain do not undergo Interkinetic Nuclear Migration (INM), as in Lissencephaly where affected children have severe psychomotor retardation^1^. In oocytes from worm, sea urchin and frog the location of the nucleus marks the animal pole and in *Drosophila* it defines the future dorso-ventral axis of the embryo and of the adult body plan^2–4^. However, in mouse and humans, the oocyte nucleus is centrally located at the end of oocyte growth in Prophase I, oocytes display no obvious sign of polarity^5,6^ and thus nuclear position does not instruct embryo axis. Nevertheless an off-centred nucleus correlates with a poor outcome of oocyte development^7,8^, arguing that central positioning is important for later embryo development. This is surprising since oocytes further undergo two asymmetric divisions in size of daughter cells, requiring an off-centring of their chromosomes^9^. It is therefore of fundamental importance to decipher the biological significance of nucleus centring in mammalian oocytes.

Mouse oocytes lack canonical centrosomes, major MicroTubule Organizing Centres (MTOCs) of cells and thus do not use centrosome-based nucleus positioning, as in many other systems like migrating neurons or *Drosophila* oocytes^1,10^. In mouse oocytes, the nucleus is positioned to the centre via actin-based mechanisms and microtubules have a minor contribution to this phenomenon^11^. Using a multidisciplinary approach combining cell biology, biophysics and signal processing methods coupled to modelling, we previously have shown how the nucleus robustly localizes in an original manner to the centre of mouse oocytes^11^. Oocytes from the Formin 2 (*Fmn2*) knockout strain, which lack microfilaments in their cytoplasm, present off-centred nuclei^12,13^. Formin 2 is a straight microfilament nucleator and also an essential maternal gene since *Fmn2* total knockout females are hypofertile^14^. The re-introduction of Formin 2 in *Fmn2*^*−/−*^ oocytes, with initially off-centred nuclei, induces the motion of the nucleus towards the centre, completed in about 5 hours, in 100 % of the cases^11^. An effective pressure gradient, mediated by Formin 2-nucleated F-actin vesicles allows nucleus centring, such that at steady state, the forces are balanced and the nucleus remains centrally located^11^. Here, we discovered that nuclear architecture is unexpectedly altered in *Fmn2*^*−/−*^ oocytes. Using a computational bio-imaging approach we extracted features that show statistically significant differences between controls and *Fmn2*^*−/−*^ oocyte nuclei. Consistent with differences in nuclear architecture, RNA sequencing of control versus *Fmn2*^*−/−*^ oocytes detected 2285 genes showing significant differential expression. Mechanical rescue of the process of nucleus positioning correlated with the rescue in nuclear architecture and rescue of some gene expression. Using biophysical modelling, we show that F-actin mediated cytoplasmic activity increases nuclear envelope fluctuations and enhances chromatin diffusion inside the nucleus. Our multidisciplinary approach proposes a comprehensive model whereby mechano-transduction via microfilaments modulates nuclear architecture and the amount of maternal transcripts, instrumental to sustain early embryonic development and thus essential for gamete fitness.

## Results

### Nuclear organisation differs in *Fmn2*^*+/−*^ versus *Fmn2*^*−/−*^ oocytes

In addition to presenting off-centred nuclei, *Fmn2*^*−/−*^ oocytes displayed different shaped nuclei than control counterparts^11^. We therefore assessed nuclear architecture in *Fmn2*^*+/−*^ versus *Fmn2*^*−/−*^ oocytes. We immunostained chromatin and Lamin A in the two types of oocytes (Fig. 1A). The nucleus of *Fmn2*^*−/−*^ oocytes presented an important smooth surface and a major invagination, not observed in controls (Fig. 1A white arrow). In *Fmn2*^*+/−*^ and *Fmn2*^*−/−*^ nuclei, the Lamin A staining was uninterrupted all along the nuclear envelope and the intensity of this staining was comparable, suggesting that the endogenous amount of Lamin A was not affected in these oocytes (Fig. 1A). To address the integrity of the nuclear envelope in *Fmn2*^*−/−*^ oocytes, we used a nucleoplasmic probe that leaks out of the nucleus when nuclear envelope permeability increases^13^. The nucleo-cytoplasmic ratio of the probe is not significantly different between *Fmn2*^*+/−*^ and *Fmn2*^*−/−*^ oocytes, arguing that the nuclear envelope is intact in *Fmn2*^*−/−*^ oocytes (Extended Data Fig. 1). To quantify the observed differences in nuclear architecture between *Fmn2*^*+/−*^ and *Fmn2*^*−/−*^ oocytes, we developed a computational imaging approach to automatically threshold the stacks of images and extract non-redundant features (shape, volume, DNA dispersion: Fig. 1A and see Supplementary Methods for the full list of features). We extracted 46 non-redundant features of nucleus architecture that differ significantly between control and *Fmn2*^*−/−*^ oocytes. To illustrate, we presented 2 of these features (Fig. 1B and C). In *Fmn2*^*−/−*^ oocytes, the DNA was more dispersed and presented a higher distance of the DNA centroid to the outer DNA surface (Fig. 1B compare green and red dots). This was not due to an increase in nuclear size (mean and SD of nuclear volume of 3485 +/− 708.3 μm^3^ for *Fmn2*^*+/−*^ and 3867 +/− 1857 μm^3^ for *Fmn2*^*−/−*^, non significant with Mann-Whitney U p-value of 0.48). Also contacts between DNA and Lamin A significantly increased in *Fmn2*^*−/−*^ oocytes compared to controls (Fig. 1C compare green and red dots). Our computational approach thus allowed to detect and to quantify differences in nuclear architecture of the two types of oocytes.

**Figure 1:**
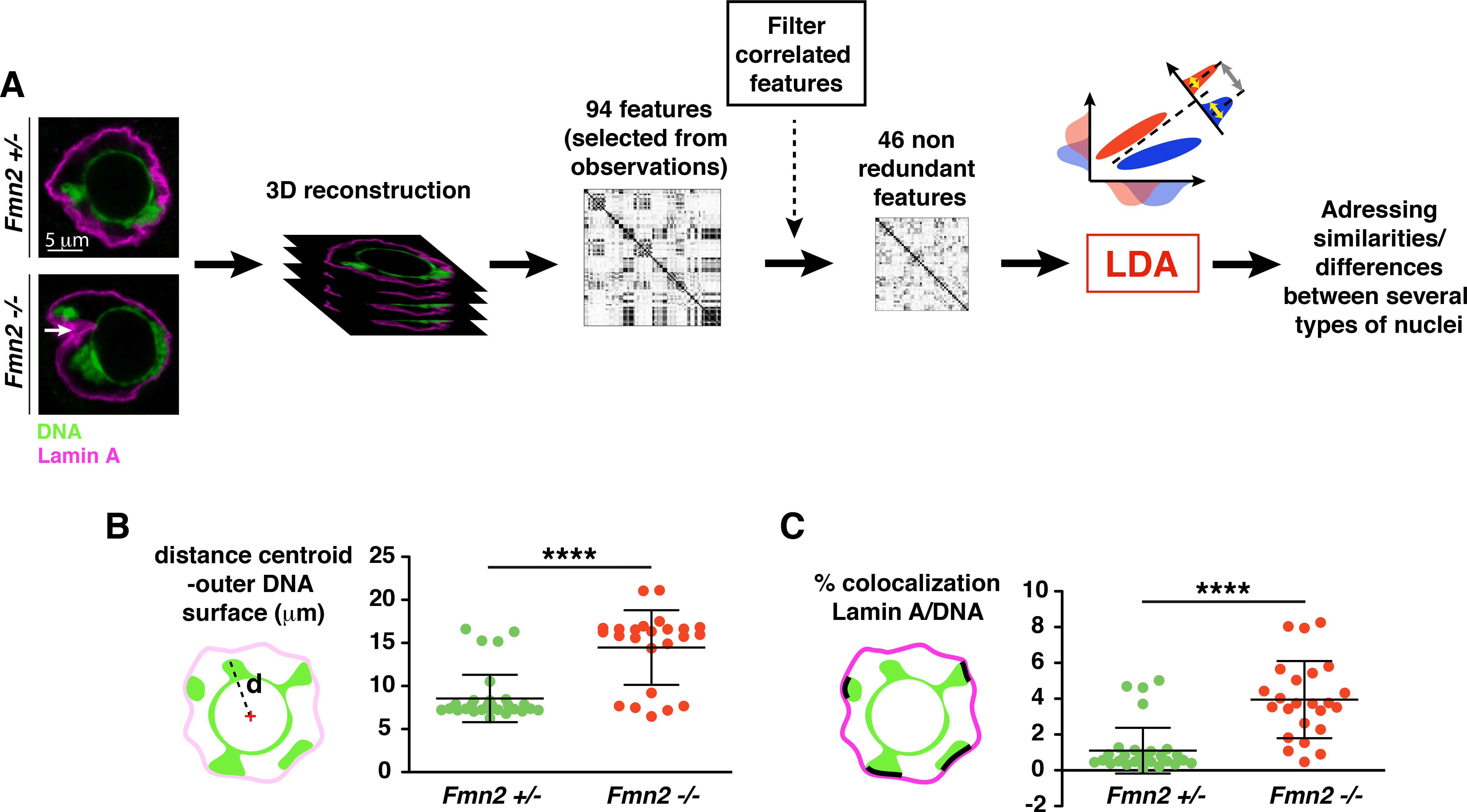
Nuclear architecture defects in absence of Formin 2 highlighted by a computational imaging approach. **A:** Principle of the computational imaging approach to analyse nuclear architecture. Oocytes were stained for Lamin A (magenta) and DNA (green), as shown on the single plane images of a *Fmn2*^*+/−*^ and a *Fmn2*^*−/−*^ nucleus (top and bottom left panels). The white arrow points to the invagination that can be observed in *Fmn2*^*−/−*^ nuclei. The thresholds are the same for both images. Scale bar is 5 μm. The nuclei were reconstructed in 3D to quantify their global architecture. 94 features describing nuclear architecture were selected from observations, such as for example shape, volume, DNA dispersion, DNA fragmentation, overlap between Lamin A and DNA, and other features connecting Lamin A to DNA (see Supplementary Methods for the full list of features). Correlated features were eliminated via a correlation matrix (see Methods) and 46 non-redundant features were eventually used for the analysis. Remaining features were then subjected to a Linear Discriminant Analysis (LDA), where the 46 features dataset is projected onto a two-dimensional space by maximazing the variance between different classes (gray double-arrow) while minimizing the variance within the same class (yellow double-arrows), providing a powerful tool to classify the different types of nuclei (see Methods for a more detailed description). **B:** Nuclear architecture is perturbed in *Fmn2*^*−/−*^ oocytes compared to *Fmn2*^*+/−*^ oocytes. Measure of the feature « DNA dispersion » extracted from the computational imaging approach. Left: Scheme of the principle of the DNA dispersion measure, which corresponds to the distance from the centroid to the outer surface of the DNA mass. Error bars represent SD. Mean and SD are 8.55 +/− 2.74 μm for *Fmn2*^*+/−*^ and 14.47 +/− 4.33 μm for *Fmn2*^*−/−*^. Mann-Whitney U p-value = 8.06E-06. N=35 oocytes for *Fmn2*^*+/−*^, 2 independent experiments and N=24 oocytes for *Fmn2*^*−/−*^, 2 independent experiments. **C:** Nuclear architecture is perturbed in *Fmn2*^*−/−*^ oocytes compared to *Fmn2*^*+/−*^ oocytes. Measure of the feature « overlap between Lamin A and DNA » extracted from the computational imaging approach. Left: Scheme of the principle of the measure of the overlap between Lamin A and DNA, which is the percentage of colocalization of Lamin A and DNA related to the total DNA. Error bars represent SD. Mean and SD are 1.1 +/− 1.28 for *Fmn2*^*+/−*^ and 3.95 +/− 2.16 for *Fmn2*^*−/−*^. Mann-Whitney U p-value = 2.55E-07. N=35 oocytes for *Fmn2*^*+/−*^, 2 independent experiments and N=24 oocytes for *Fmn2*^*−/−*^, 2 independent experiments.

### The transcriptome of *Fmn2*^*+/−*^ and *Fmn2*^*−/−*^ oocytes are different

A-type Lamins are known to regulate gene expression^15,16^. We therefore compared the transcriptome of *Fmn2*^*+/−*^ to the one of *Fmn2*^*−/−*^ oocytes using RNA sequencing. We found that 2285 genes, out of 23474 (about 10% of analysed genes), were significantly (padj < 0.05) mis-regulated between the two types of oocytes. Amongst these, 128 were under-expressed while 65 were over-expressed more than 2 fold in *Fmn2*^*−/−*^ oocytes (Fig. 2A). We validated 10 of these deregulated genes by RTqPCR (Fig. 2C and Extended Data Table 1). A gene-ontology analysis of the 2285 deregulated genes showed a specific enrichment in processes involved in meiotic divisions and early development, such as for example Tcstv1, a gene involved in Zygotic genome activation (Fig. 2B and Extended Data Table 1). The RNA sequencing data argued that changes in nuclear architecture in *Fmn2*^*−/−*^ oocytes correlates with modification in gene expression.

**Figure 2:**
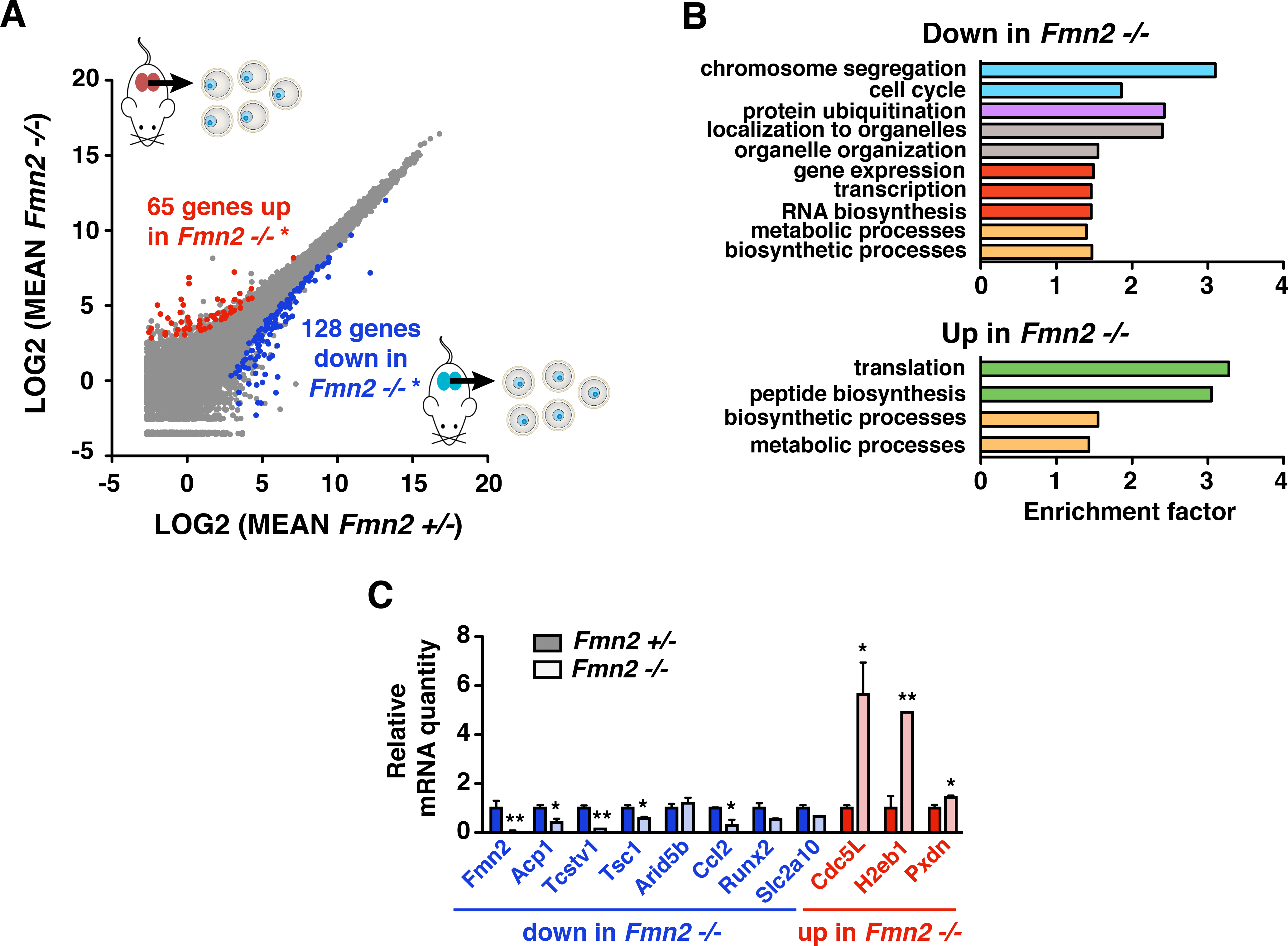
Gene expression is altered in *Fmn2*^*−/−*^ Prophase I oocytes compared to controls. **A:** Differential gene expression in *Fmn2*^*−/−*^ Prophase I-arrested oocytes. Differential transcriptomic analysis of *Fmn2*^*+/−*^ vs *Fmn2*^*−/−*^ fully-grown oocytes performed by RNA sequencing. Among the 23474 detected transcripts, 128 are significantly downregulated and 65 upregulated more than 2-fold in *Fmn2*^*−/−*^ oocytes (adjusted p-value padj < 0.05 and | log2 fold change | > 1). For each condition *Fmn2*^*+/−*^ and *Fmn2*^*−/−*^, 2 biological replicates of 50 oocytes each and 3 technical replicates. **B:** Differentially expressed genes in *Fmn2*^*−/−*^ have important functions in meiotic and first embryonic divisions. Gene Ontology enrichment analysis for Biological Processes performed on the genes expressed differentially in *Fmn2*^*−/−*^ oocytes with an adjusted p-value padj < 0.05: 1365 downregulated (top graph) and 920 upregulated (bottom graph), from RNAseq data. Enrichment factor corresponds to the representation of a category of Biological Processes within the analysed gene database compared to a mouse reference database. P-values for PANTHER over-representation test with Bonferroni correction are all < 0.05. **C:** Confirmation of the transcriptomics analysis performed in A by RT-qPCR on a panel of 8 downregulated and 3 upregulated genes selected from their adjusted p-value and | log2 fold change |. Mean of the normalized ratios of transcript levels between *Fmn2*^*+/−*^ and *Fmn2*^*−/−*^. Error bars represent normalized SEM. Normalized mean and SEM for *Fmn2*^*+/−*^ are 1 +/− 0.29 (Fmn2), 1 +/− 0.13 (Acp1), 1 +/− 0.11 (Tcstv1), 1 +/− 0.11 (Tsc1), 1 +/− 0.18 (Arid5b), 1 +/− 0.027 (Ccl2), 1 +/− 0.2 (Runx2), 1 +/− 0.12 (Slc2a10), 1 +/− 0.11 (Cdc5L), 1 +/− 0.49 (H2eb1) and 1 +/− 0.13 (Pxdn). Normalized mean and SEM for *Fmn2*^*−/−*^ are 0.065 +/− 0.02 (Fmn2), 0.41 +/− 0.15 (Acp1), 0.15 +/− 0.006 (Tcstv1), 0.58 +/− 0.068 (Tsc1), 1.2 +/− 0.22 (Arid5b), 0.3 +/− 0.23 (Ccl2), 0.54 +/− 0.036 (Runx2), 0.66 +/− 0.021 (Slc2a10), 5.64 +/− 1.3 (Cdc5L), 4.9 +/− 0.016 (H2eb1) and 1.44 +/− 0.08 (Pxdn). P-values for Student t test are 0.01 for Fmn2, 0.043 for Acp1, 0.0014 for Tcstv1, 0.034 for Tsc1, 0.53 for Arid5b, 0.022 for Ccl2, 0.084 for Runx2, 0.051 for Slc2a10, 0.012 for Cdc5L, 0.0013 for H2eb1 and 0.048 for Pxdn. 3 independent experiments, N=35 oocytes each for Acp1, Tcstv1, Tsc1, Arid5b, Runx2, Slc2a10, H2eb1 and Pxdn, and 4 experiments, N=35 oocytes each for Fmn2, Ccl2, and Cdc5L.

### Expressing full length Formin 2 in *Fmn2*^*−/−*^ oocytes rescues nuclear position and nuclear architecture

We then tested whether nuclear position correlates with nuclear architecture and defects in gene expression observed in *Fmn2*^*−/−*^ oocytes. We injected into these oocytes cRNAs encoding either full length Formin 2, or only the actin-nucleating domain of Formin 2, the FH1-FH2 domain^17^, or the N-terminal part of Formin 2, which does not nucleate actin (Fig. 3A). Only the full length^11^ and the FH1-FH2 domain of Formin 2 were able to nucleate a cytoplasmic actin meshwork and rescue nucleus central position. The N-terminal domain of Formin 2 had no effect on nucleus position inside the oocyte and thus was probably not able to nucleate actin. We further compared nuclear architecture in the three conditions (Fig. 3A) with that of *Fmn2*^*−/−*^ oocytes using our dedicated computational 3D imaging approach. Both full length and FH1-FH2 restored the spherical shape of nuclei, with no major invagination, as in controls (Fig. 3B). Both also restored the percentage of contacts with Lamin A (Fig. 3D; right panel). The full length Formin 2 could significantly rescue the spreading of DNA inside the nucleus (Fig. 3D left panel). The FH1-FH2 was less efficient for this feature (Fig. 3D left panel) suggesting that the FH1-FH2 domain does not fully recapitulate all endogenous interactions of Formin 2. At that stage, in Prophase I, there is no strict correlation between chromatin condensation and gene expression^18^. Thus the contact between Lamin A and DNA (Fig. 3D right panel) could be more relevant to nuclear architecture than chromatin condensation. The perfect rescue of nuclear architecture by full length Formin 2 was also attested by our Linear Discriminant Analysis (LDA, see Fig. 1A) where the category of *Fmn2*^*+/−*^ and (*Fmn2*^*−/−*^ + Fmn2) nuclei could not be dissociated (Fig. 3C, compare the distribution of green versus blue dots). This category was separated from the one of *Fmn2*^*−/−*^, which was further apart on the X axis, where most of the variance (72%) lies (Fig. 3C distribution of green versus red dots), attesting that nuclear architecture differs between controls and *Fmn2*^*−/−*^ oocytes. Our LDA analysis confirmed that the FH1-FH2 domain did not rescue all features of nuclear organisation present in *Fmn2*^*−/−*^ (Fig. 3C, compare the distribution of purple to that of green dots). Yet, rescue with full-length and with the FH1-FH2 overlap when projected on the X axis, which corresponds to most of the variance (72%), attesting that the truncated form of Formin 2 rescues some aspects of nuclear organisation (Fig. 3C). Nuclear architecture can be efficiently rescued by expression of the full-length Formin 2 in *Fmn2*^*−/−*^ oocytes supporting the hypothesis that the process of nucleus centring impacts nuclear architecture.

**Figure 3:**
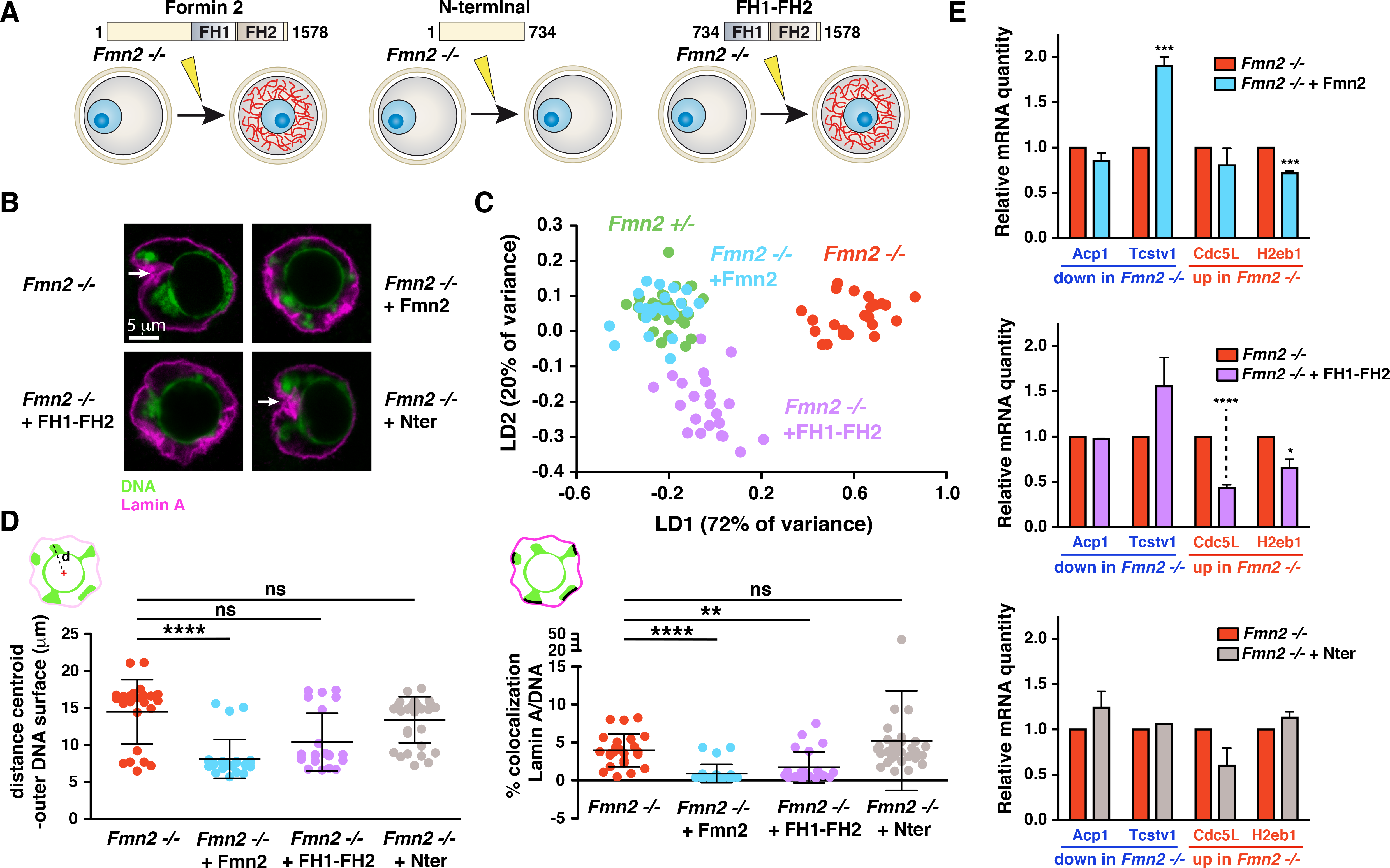
Nuclear architecture and gene expression can be rescued by full length Formin 2 expression. **A:** Scheme of the different rescue experiments testing for a role of nucleus centring in the regulation of nuclear architecture and gene expression. Left: *Fmn2*^*−/−*^ oocytes are injected with cRNAs coding for full-length Formin 2, leading to the reformation of a cytoplasmic actin meshwork (red filaments), and incubated for 8 hours to allow nucleus centring. Middle: *Fmn2*^*−/−*^ oocytes are injected with cRNAs coding for the N-terminal domain of Formin 2 and incubated for 8 hours. No meshwork reformation and no nucleus centring are observed in these conditions. Right: *Fmn2*^*−/−*^ oocytes are injected with cRNAs coding for the FH1-FH2 actin-nucleating domain of Formin 2, leading to the reformation of a cytoplasmic actin meshwork (red filaments), and incubated for 8 hours to allow nucleus centring. The size of full-length Formin 2 is 1578 amino acids (aa), Formin 2 N-terminal domain corresponds to aa 1 to 734 and Formin 2 FH1-FH2 domain corresponds to aa 734 to 1578. **B:** Nuclear architecture in *Fmn2*^*−/−*^ and in the 3 different rescue conditions. Single plane images of nuclei stained for Lamin A (magenta) and DNA (green) in *Fmn2*^*−/−*^ (top left), *Fmn2*^*−/−*^ expressing either the full-length Formin 2 (Fmn2, top right), the FH1-FH2 domain (FH1-FH2, bottom left) or the N-terminal domain (Nter, bottom right). White arrows point to the invaginations than can be observed in nuclei from *Fmn2*^*−/−*^ and *Fmn2*^*−/−*^ expressing the N-terminal domain. Thresholding is the same for all images. Scale bar is 5 μm. **C:** Nuclear architecture is regulated by nucleus centring. Linear Discriminant Analysis (LDA) of the 4 different types of nuclei *Fmn2*^*+/−*^, *Fmn2*^*−/−*^, *Fmn2*^*−/−*^ + full-length Formin 2 and *Fmn2*^*−/−*^ + FH1-FH2 domain (see Fig. 1A and Methods for a more detailed explanation). The first axis (LD1) of the two-dimensional space, where the 46 features dataset for each oocyte is projected, represents 72% of the variance whereas the second axis (LD2) represents 20% of the variance. N=35 for *Fmn2*^*+/−*^, 2 independent experiments, N=24 for *Fmn2*^*−/−*^, 2 independent experiments, N=26 for *Fmn2*^*−/−*^ + Fmn2, one experiment, N=23 for *Fmn2*^*−/−*^ + FH1-FH2, one experiment. **D:** Only full length Formin 2 and the FH1-FH2 domain of Formin 2 can rescue the co-localization of DNA with Lamin A. Measure of the extent of rescue of nuclear architecture of *Fmn2*^*−/−*^ oocytes by full-length Formin 2, by the FH1-FH2 domain and by the N-terminal domain as represented by the 2 features « DNA dispersion » and « overlap between Lamin A and DNA » from the computational imaging approach described in Fig. 1A. Left: DNA dispersion, measured by the distance from the centroid to the outer surface of the DNA mass. Error bars represent SD. Mean and SD are 14.47 +/− 4.33 μm for *Fmn2*^*−/−*^, 8.09 +/− 2.64 μm for *Fmn2*^*−/−*^ + Fmn2, 10.36 +/− 3.90 μm for *Fmn2*^*−/−*^ + FH1-FH2 and 13.38 +/− 3.13 μm for *Fmn2*^*−/−*^ + Nter. Mann-Whitney U p-values are 7.2E-06 for *Fmn2*^*−/−*^ - *Fmn2*^*−/−*^ + Fmn2, 3.33E-02 for *Fmn2*^*−/−*^ - *Fmn2*^*−/−*^ + FH1-FH2 and 4.62E-02 for *Fmn2*^*−/−*^ - *Fmn2*^*−/−*^ + Nter. N=24 for *Fmn2*^*−/−*^, 2 independent experiments, N=26 for *Fmn2*^*−/−*^ + Fmn2, one experiment, N=23 for *Fmn2*^*−/−*^ + FH1-FH2, one experiment and N= 32 for *Fmn2*^*−/−*^ + Nter, 2 independent experiments. Right: overlap between Lamin A and DNA, measured by the percentage of co-localization of Lamin A and DAPI related to the total DNA. Error bars represent SD. Mean and SD are 3.95 +/− 2.16 for *Fmn2*^*−/−*^, 0.91 +/− 1.19 for *Fmn2*^*−/−*^ + Fmn2, 1.74 +/− 2.05 for *Fmn2*^*−/−*^ + FH1-FH2 and 5.24 +/− 6.56 for *Fmn2*^*−/−*^ + Nter. Mann-Whitney U p-values are 8.58E-07 for *Fmn2*^*−/−*^ - *Fmn2*^*−/−*^ + Fmn2, 8.16E-04 for *Fmn2*^*−/−*^ - *Fmn2*^*−/−*^ + FH1-FH2 and 8.01E-01 for *Fmn2*^*−/−*^ - *Fmn2*^*−/−*^ + Nter. N=24 for *Fmn2*^*−/−*^, 2 independent experiments, N=26 for *Fmn2*^*−/−*^ + Fmn2, one experiment, N=23 for *Fmn2*^*−/−*^ + FH1-FH2, one experiment and N= 32 for *Fmn2*^*−/−*^ + Nter, 2 independent experiments. **E:** Only the full length Formin 2 and the FH1-FH2 domain, but not the N-terminal domain can rescue gene expression defects from *Fmn2*^*−/−*^ oocytes. Rescue was addressed by RT-qPCR on 4 differentially expressed genes in *Fmn2*^*−/−*^ confirmed in Fig. 2C (2 downregulated and 2 upregulated). Top: Mean of the normalized ratios of transcript levels between *Fmn2*^*−/−*^ and *Fmn2*^*−/−*^ + Fmn2. Error bars represent SEM. Mean and SEM are 0.85 +/− 0.090 (Acp1), 1.90 +/− 0.10 (Tcstv1), 0.80 +/− 0.19 (Cdc5L), 0.72 +/− 0.029 (H2eb1). P-values for t-test are 0.1698 (Acp1), 0.0008 (Tcstv1), 0.355 (Cdc5L) and 0.0006 (H2eb1). 3 independent experiments, N=20 oocytes by condition and by experiment. Middle: Mean of the normalized ratios of transcript levels between *Fmn2*^*−/−*^ and *Fmn2*^*−/−*^ + FH1-FH2. Error bars represent SEM. Mean and SEM are 0.97 +/− 0.009 (Acp1), 1.56 +/− 0.32 (Tcstv1), 0.44 +/− 0.033 (Cdc5L), 0.65 +/− 0.097 (H2eb1). P-values for t-test are 0.058 (Acp1), 0.1567 (Tcstv1), <0.0001 (Cdc5L) and 0.0119 (H2eb1). 4 independent experiments, N=20 oocytes by condition and by experiment. Bottom: Mean of the normalized ratios of transcript levels between *Fmn2*^*−/−*^ and *Fmn2*^*−/−*^ + Nter. Error bars represent SEM. Mean and SEM are 1.24 +/− 0.18 (Acp1), 1.06 +/− 0.002 (Tcstv1), 0.60 +/− 0.19 (Cdc5L), 1.13 +/− 0.064 (H2eb1). P-values for t-test are 0.2483 (Acp1), 0.1194 (Tcstv1), 0.1086 (Cdc5L) and 0.111 (H2eb1). 3 independent experiments, N=20 oocytes by condition and by experiment.

### Expressing full length Formin 2 in *Fmn2*^*−/−*^ oocytes rescues gene expression

Since we could rescue nuclear positioning as well as to some extent nuclear organisation in a few hours using both full length and FH1-FH2 domain of Fmn2, we tested several candidates for gene expression rescue. Full length Formin 2 rescued both down as well as up-regulated genes from *Fmn2*^*−/−*^ oocytes. It significantly rescued two out of the four tested genes (Fig. 3E upper panel). The rescue of two genes by full length Fmn2, Tcstv1 and H2eb1 was most certainly due to an effect on transcription since it was abolished by simultaneous treatment with RNA polymerase II inhibitor α-amanitin during the nucleus repositioning assay^11^ (Extended Data Fig. 2). As known for decades but never re-addressed recently with more performing tools, mouse oocyte growth terminates with global transcription silencing^19–21^. We confirmed this observation by EU incorporation in incompetent (with non surrounded nucleolus) versus competent (with surrounded nucleolus) oocytes (Extended Data Fig. 3A). Incompetent oocytes, which have not finished their growth, was approximately 5 times more transcriptionally active than competent fully-grown ones (Extended Data Fig. 3B). Surprisingly, we consistently detected few foci of EU incorporation in fully-grown competent oocytes (Extended Data Fig. 3A and B). This novel observation suggests that even if, as observed decades ago, there is indeed a global shut down of transcription during the end of oocyte growth, few loci are still active in fully grown oocytes prior to meiosis resumption and can be re-activated in our rescue experiments. We can draw a parallel with recent findings in *Drosophila* oocytes, where, consistent with older results, transcriptional reactivation has been observed in stage 9 Prophase I arrested oocytes^22,23^, against the commonly accepted idea of transcriptional shut down during the full Prophase I arrest. Residual transcription has been found even in mitosis, favouring transcriptional reactivation during mitotic exit^24^.

It is noteworthy that the gene Tcstv1, which is the shortest (1399 bp compared to 18287 bp for Acp1), is the one most significantly rescued in this 8 hours assay. This might be related to the processivity of the Pol II polymerase about 30 nt s^−1^ ^25^. Indeed, the response of our model system seems rather slow. Acute disruption of the cytoplasmic mesh using Cytochalasin D (CCD, an actin depolymerising drug) on control oocytes leads to a significant up-regulation of Cdc5L (3.99 +/− 0.54 fold) comparable to its up-regulation in *Fmn2*^*−/−*^ oocytes (5.64 +/− 1.3 fold) only after 5 hours (Extended data Fig. 3C). The FH1-FH2 construct rescued two genes out of four (Fig. 3E middle panel). Importantly, the N-terminal construct, that does not nucleate actin, had no impact on nucleus positioning, on nuclear shape (Fig. 3B), on chromatin spreading (Fig. 3D left panel), on the extent of contacts between Lamin A and chromatin (Fig. 3D right panel) and did not rescue any of the four genes that were tested (Fig. 3E lower panel). Altogether, our results suggest that F-actin dependent nucleus centring impacts nuclear architecture as well as gene expression in mouse oocytes.

### *Fmn2*^*−/−*^ oocytes display similar levels of expression of MAL-SRF responding genes and similar levels of nuclear G-actin than *Fmn2*^*+/−*^ oocytes

What is the mechanism potentially responsible for this correlation between F-actin dependent nucleus centring and gene expression? Direct regulation of gene expression by nuclear G-actin is well documented (for a review see ^26^). One of the best described systems is the MAL-SRF pathway^27^. In this serum-inducible system, a Serum Response Factor (SRF) requires its co-activator MAL to achieve transcription of target genes. MAL translocates to nuclei, but its binding to monomeric nuclear actin favors its export to the cytoplasm, thus preventing transcriptional activation. Upon serum stimulation, polymerization of cytoplasmic F-actin decreases globally the monomeric G-actin pool, thus increasing the nuclear amount of MAL free from G-actin, which induces transcription of target genes^27^.

The responsiveness of the MAL/SRF pathway is shorter than the one provided by nucleus positioning: 45 to 90 min for SRF target genes^27^, compared to 5 to 8 hours in our model system where nuclei are the size of a somatic cell (30 μm wide). This may suggest different underlying mechanisms. Furthermore, from our RNAseq data, we checked whether important players of the MAL-SRF pathway were transcriptionally deregulated in oocytes lacking Formin 2: the SRF transcription factor, its MAL co-activator, and also MICAL-2, an upstream regulator of the MAL/SRF pathway^28^, as well as MAL-dependent SRF target genes *Vcl*, *Srf*, *Cyr61*^27^ None of these genes were deregulated in the absence of Formin 2 in our RNA seq data.

Nevertheless, we addressed the potential contribution of nuclear actin with MAL as an actin monomer sensor. We used the RPEL1-eGFP-3NLS probe^29^, where RPEL1, the actin monomer-binding domain of MAL is fused to eGFP and targeted to the nucleus by 3 Nuclear Localization Sequences (Fig. 4A). This probe also indirectly reports for the MAL-SRF activation status. As expected, the localization of this probe was nuclear (Fig. 4B). The amount of RPEL1-eGFP-3NLS was comparable between *Fmn2*^*+/−*^ and *Fmn2*^*−/−*^ nuclei (Fig. 4C). As a positive control of the sensitivity of the probe, its amount was increased in *Fmn2*^*+/−*^ nuclei after a 3h treatment with CCD (Fig. 4C). Despite this, Cdc5L was not yet up-regulated at 3h by CCD treatment (Extended data Fig. 3C and Fig. 4C), which suggested that transcriptional regulation is not impacted by the level of nuclear actin in oocytes at that stage. Our results are consistent with RPEL1 as a good nuclear actin monomer sensor and show that deregulation of gene expression in *Fmn2*^*−/−*^ oocytes cannot be explained by changes in nuclear actin nor activation of the MAL-SRF pathway.

**Figure 4:**
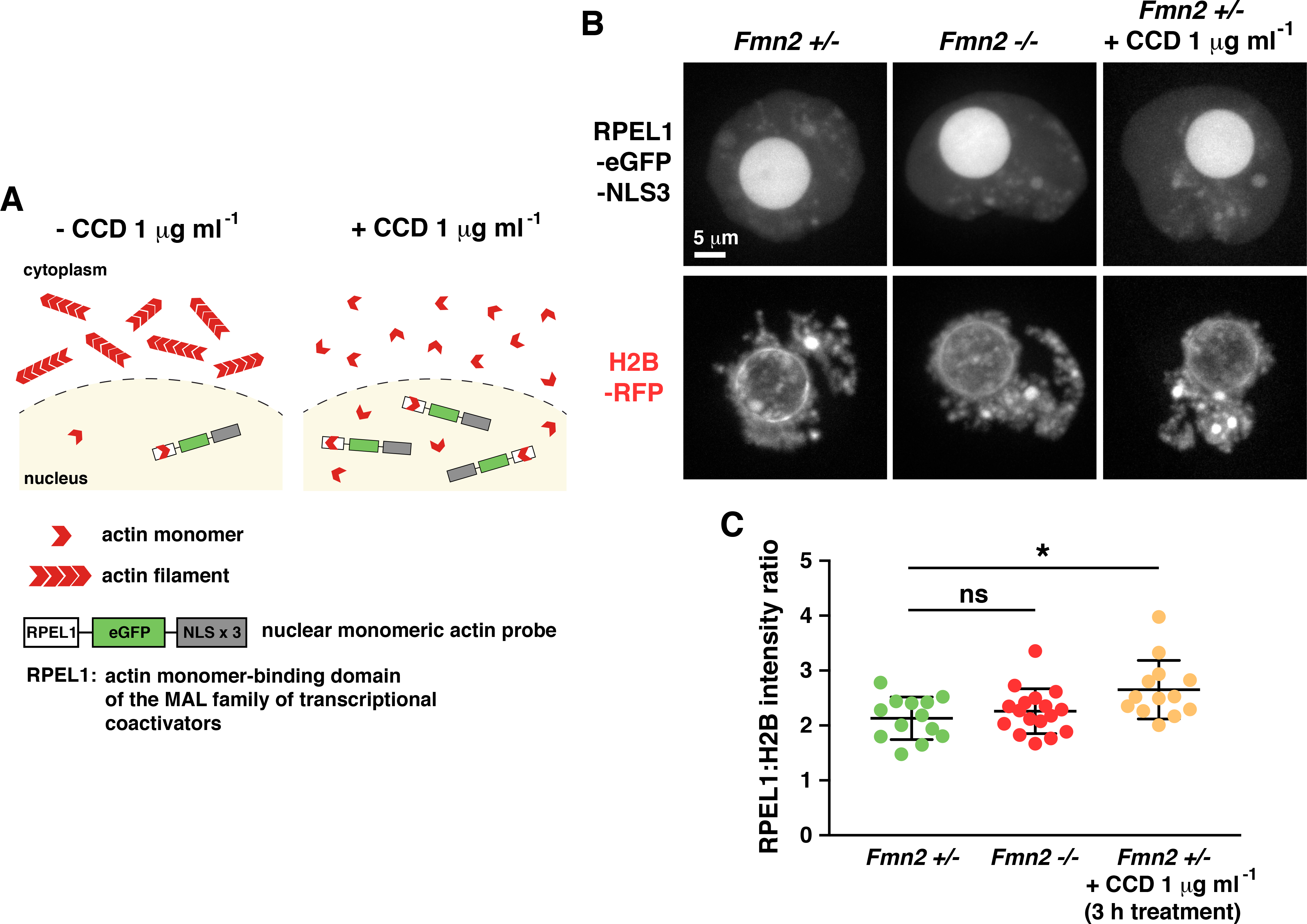
*Fmn2*^*+/−*^ and *Fmn2*^*−/−*^ nuclei present similar levels of nuclear actin. **A:** Principle of the RPEL1-eGFP-NLS3 nuclear monomeric actin probe. The probe is a fusion of RPEL1, an actin monomer-binding domain of the MAL family of transcriptional coactivators, eGFP and 3 nuclear localization sequences. The probe binds to monomeric actin in the nucleus through the RPEL1 domain. Without CCD, a large part of the actin pool is incorporated within the cytoplasmic actin meshwork. By depolymerizing the cytoplasmic actin meshwork, treatment with CCD increases the pool of nuclear monomeric actin that binds to the RPEL1-eGFP-NL3 probe (see significant increase of RPEL1 signal between *Fmn2*^*+/−*^ versus *Fmn2*^*+/−*^ + CCD 1 μg ml^−1^ nuclei in C). **B:** RPEL1-eGFP-NLS3 signal in *Fmn2*^*+/−*^ and *Fmn2*^*−/−*^ nuclei, and in *Fmn2*^*+/−*^ nuclei after 3 hours treatment with 1 μg ml^−1^ CCD. Top: RPEL1-eGFP-NLS3 signal. Bottom: H2B-RFP signal. Maximal projections of signals are presented. Scale bar is 5 μm. The fluorescence levels do not represent the intensities measured in C. **C:** Quantification of the RPEL1 signal in *Fmn2*^*+/−*^ compared to *Fmn2*^*−/−*^ and *Fmn2*^*+/−*^ + CCD oocytes. RPEL1 signal is normalized to H2B-RFP. Error bars represent SD. Mean and SD are 2.13 +/− 0.39 for *Fmn2*^*+/−*^, 2.26 +/− 0.41 for *Fmn2*^*−/−*^ and 2.65 +/− 0.53 for *Fmn2*^*+/−*^ + CCD. Mann-Whitney U p-values are 0.59 for *Fmn2*^*+/−*^ - *Fmn2*^*−/−*^ and 0.01 for *Fmn2*^*+/−*^ - *Fmn2*^*+/−*^ + CCD. N=13 for *Fmn2*^*+/−*^, 2 independent experiments, N=17 for *Fmn2*^*−/−*^, 2 independent experiments, and N=13 for *Fmn2*^*+/−*^ + CCD, 2 independent experiments.

### Actin promotes extensive nuclear envelope fluctuations

Since we excluded the contribution of two major players (nuclear actin and MAL-SRF pathway) in deregulation of gene expression in *Fmn2*^*−/−*^ oocytes, we checked for a potential mechano-transduction effect. Indeed the 5h-CCD treatment of *Fmn2*^*+/−*^ oocytes, which maintain centrally located nuclei due to the high viscosity of the cytoplasm^11^, affected to a similar extent the expression of a highly up-regulated gene in *Fmn2*^*−/−*^ (Extended data Fig. 3C). This observation is in favour of F-actin mediated forces acting at steady state on the nuclear envelope. Labelling the nucleus using a nucleoplasmic probe^13^ allows to follow the nuclear envelope contour at high temporal resolution (stream mode) in 2D movies and to follow nuclear deformations at steady state. As observed before^30^, the nuclear envelope of mouse oocytes in Prophase I is highly deformable (Fig. 5A upper panel left and Supplementary Video 1). The nucleus from a control oocyte (*Fmn2*^*+/−*^) was more or less round with deformations of its nuclear envelope evenly distributed (Fig. 5A). In contrast, but consistent with our observations on fixed cells (Fig. 1A), nuclei from *Fmn2*^*−/−*^ appear deformed, no longer round and presenting 2/3 (65.76%) of their surface smooth (Fig. 5A lower panel left and Supplementary Video 2). To quantify the evolution of the nuclear shape as a function of time, we measured its radius r from the centre of mass at each time point in all directions in the focal plane (i.e.: for all points of the nuclear circumference scanned using a revolving angle θ of 1° increment from 0 to 360°). The measure of the mean radius r over time for each value of θ, denoted R, allows to extract the mean 2-dimensional shape (Fig. 5A right panels) as well as the fluctuations represented by the variance (r-R)^2^ of the nuclear shape (Fig. 5B and C and see Methods). The measures showed that *Fmn2*^*+/−*^ have more regular and round shaped nuclei than *Fmn2*^*−/−*^ and that their nuclear envelope is subjected to more important fluctuations distributed throughout their entire surface (compare heat maps of Fig. 5A right from upper versus lower panels). The nuclear envelope of *Fmn2*^*+/−*^ oocytes is subjected to fluctuations whose variance is 6 times higher than in *Fmn2*^*−/−*^ oocytes (Fig. 5C). These fluctuations can be attributed to the presence of actin filaments in the cytoplasm, since 1 h 30 treatment of *Fmn2*^*+/−*^ oocytes with CCD mimicked both the nuclear shape (Extended Data Fig. 4A left with 66.12% of smooth surface), the distribution and the amount of fluctuations present in *Fmn2*^*−/−*^ oocytes (Extended Data Fig. 4A right panel and 4B-C, Supplementary Video 3). Hence, Formin 2 nucleated actin filaments promote important nuclear envelope deformations in control oocytes.

**Figure 5:**
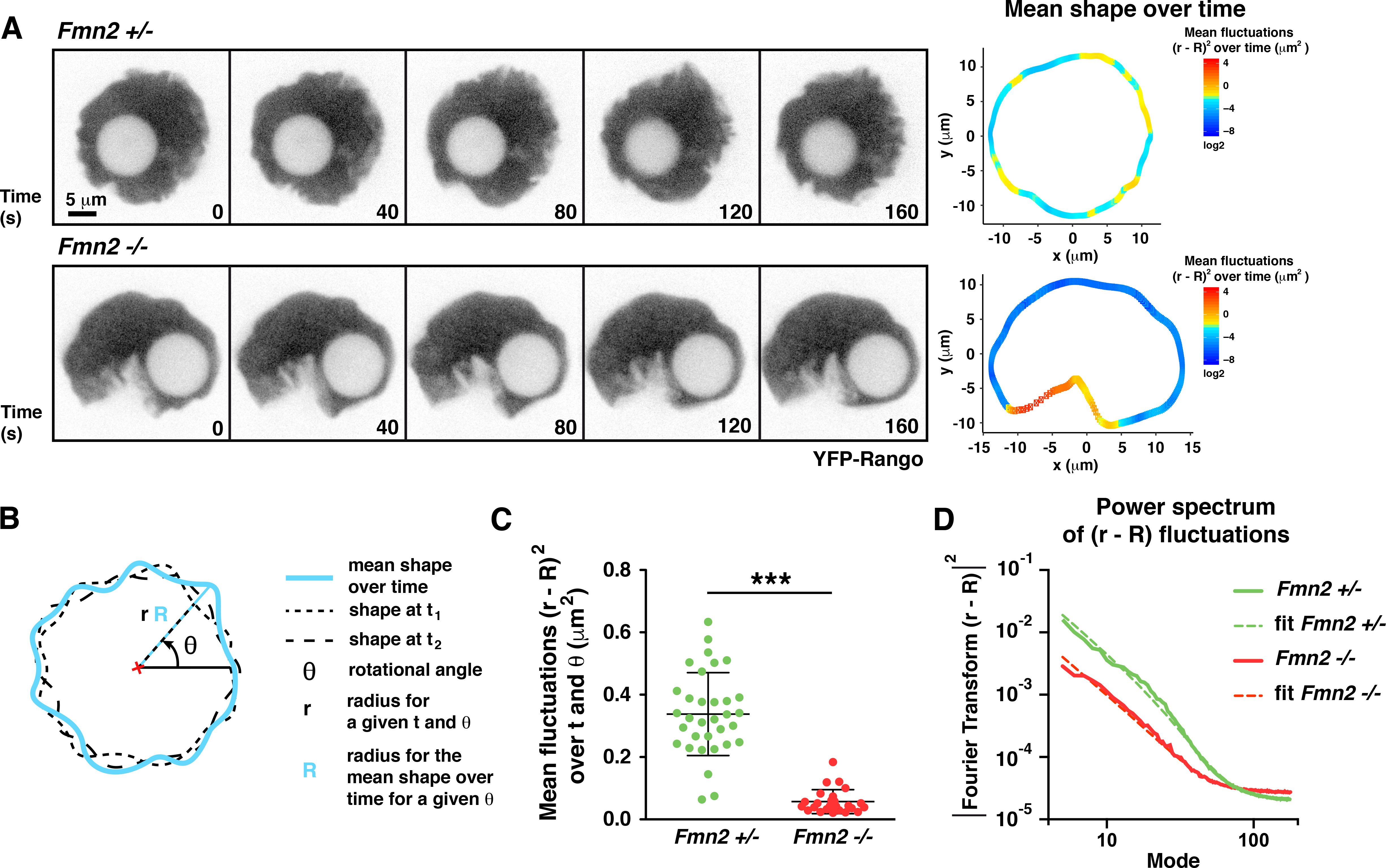
Cytoplasmic activity induces nuclear envelope fluctuations. **A:** Cytoplasmic activity impacts on nuclear envelope fluctuations and nuclear shape. Top left: images from a movie of a *Fmn2*^*+/−*^ oocyte injected with cRNAs coding for the nuclear probe YFP-Rango. Top right: Heat map of the mean nuclear outline over time (5 minutes) measured from the *Fmn2*^*+/−*^ oocyte presented on the left galery. Bottom left: images from a movie of a *Fmn2*^*−/−*^ oocyte injected with cRNAs coding for the nuclear probe YFP-Rango. Bottom right: Heat map of the mean nuclear outline over time (5 minutes) measured from the *Fmn2*^*−/−*^ oocyte presented on the left galery. The colour codes of both heat maps represent the nuclear outline fluctuations over time (5 minutes), in μm^2^, for each position along the circumference relative to the mean shape over time. Red is for higher values of fluctuations and blue for lower values of fluctuations. **B:** Scheme of the method used to quantify the nuclear envelope fluctuations over time and in a given direction. Directions were defined by a revolving angle θ of a 1° increment from 0 to 360°. Mean and variance of the radius distribution in a given direction were computed for each direction and reported as a schematic plot (heatmaps on Fig. 5A right) where the radius of the membrane represents the mean and the color on the membrane represents the corresponding variance. **C:** Cytoplasmic activity in controls promotes nuclear envelope fluctuations 6-fold higher than in *Fmn2*^*−/−*^ oocytes. Nuclear envelope fluctuations over 4 to 5 minutes represented by the variance of the radius over t and θ from *Fmn2*^*+/−*^ and *Fmn2*^*−/−*^ nuclei. For *Fmn2*^*−/−*^, the region of the large cleft was excluded from the measurements. Error bars represent SD. Mean and SD 0.34 +/− 0.13 μm^2^ for *Fmn2*^*+/−*^ and 0.057 +/−0.038 μm^2^ for *Fmn2*^*−/−*^. Mann-Whitney U p-value < 0.0001. N=33 oocytes for *Fmn2*^*+/−*^, 3 experiments and N=25 oocytes for *Fmn2*^*−/−*^, 3 experiments. **D:** Nuclear envelope fluctuations are generated by the cytoplasmic activity, independently of the nuclear envelope physical properties. Power spectrum of nuclear fluctuations over 4 to 5 minutes: the squared moduli of the Fourier transforms of fluctuations (r-R) are plotted as a function of the mode for *Fmn2*^*+/−*^ (green) and *Fmn2*^*−/−*^ (red) nuclei. For *Fmn2*^*−/−*^, the region of the large cleft was excluded from the measurements. Data are fitted to the function 1 / (σ*n*^*2*^ + κ*n*^4^) + Yinf according to the Helfrich model.

### Microtubules dampen nuclear envelope fluctuations

To investigate the contribution of microtubules to nuclear envelope fluctuations at steady state, *Fmn2*^*+/−*^ oocytes were treated with Nocodazole (NZ), a microtubule depolymerising drug, for 1h30 to 2 hours. It increased the fluctuations at the nuclear envelope (compare heat maps of Extended Data Fig. 5B right panels and Supplementary Video 5). The variance of these fluctuations was twice higher in Nocodazole treated oocytes than in control *Fmn2*^*+/−*^ oocytes (Extended Data Fig. 5C). These results are consistent with a stabilizing role of microtubules at the nuclear envelope. This could partly explain our former findings of a less directional nuclear trajectory in NZ treated oocytes during nuclear centration^11^.

We suspected that the invagination present in *Fmn2*^*−/−*^ oocytes was due to the presence of microtubules. Indeed, the major acentriolar MicroTubule Organizing Centre (aMTOC), normally apposed to the nucleus at that developmental stage^30^, was present close to the invagination (Extended Data Fig. 5A, Supplementary Video 4).

Treatment of *Fmn2*^*−/−*^ oocytes with Nocodazole removed the major invagination present in their nuclei and kept the whole nuclear envelope free of fluctuations, confirming the stabilizing role of microtubules at the nuclear envelope (Extended Data Fig. 5B-C lower panels and Supplementary Video 6). Thus the invagination is formed probably by the tethering of microtubules emanating from this major aMTOC to the nuclear envelope. Microtubules function here opposite to microfilaments, having a dampening effect on nuclear envelope fluctuations.

### A mathematical model describing nuclear envelope fluctuations

To analyse more precisely the fluctuations of the nuclear envelope and assess the physical properties of the envelope itself, we took a modelling approach. The oocyte cytoplasm can be modelled as an active fluid^11^. Here the term “active” refers to non-equilibrium processes (i.e. ATP dependent) involving F-actin, such as polymerisation/depolymerisation of actin filaments or actin/Myosin V interactions. The cytoplasm can be described as an active bath of effective temperature *T*_*e (cytoplasm)*_ = *T*+*T*_*a*_, where T is the usual temperature and *T*_*a*_ accounts for the fluctuations of the F-actin network. The rheology of the nuclear envelope being largely unknown, we adopt the classical Helfrich model of membranes^31^, parameterized by a tension σ and a bending modulus κ (see Supplementary model for more details). In order to confront the model to our data, we consider the radial deformations of the envelope in 2D, and parameterize the envelope by its local radial deformation h(θ,t)=r(θ,t)−R(θ), where R(θ) is the average over time of the radial coordinate r(θ,t) of the envelope at the polar angle θ (see Supplementary model). The fluctuations of the envelope are then conveniently analysed by introducing the Fourier decomposition 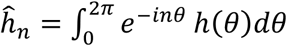, which quantifies the contribution of each mode *n*. Making use of the fluctuation dissipation theorem^32^, the power spectrum of fluctuations is then deduced and can be written (see Supplementary model):

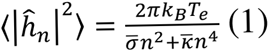

where *k*_*B*_ is the Boltzmann constant, 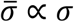 is the effective tension and 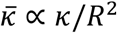 the effective bending modulus after 2D projection, with *R* the mean nucleus radius. Thus the model predicts that nuclear envelope fluctuations (quantified by the power spectrum) are controlled by the activity of the cytoplasm, which is here quantified by the active temperature *T*_*a*_. Our experimental data can be well described by a classical Helfrich model since the curves of the power spectrum as a function of the mode can be mathematically fitted with equation (1) (Fig. 5D, Extended Data Fig. 4C, Extended Data Fig. 5D). The fitting procedure gives access in particular to the characteristic length 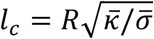, which is a marker of the physical properties of the nuclear envelope (note that the determination of tension and bending modulus independently is not accessible by this approach). Importantly, our analysis reveals that *l*_*c*_ took close values (in the range 0.24-0.4 microns) in *Fmn2*^*+/−*^, *Fmn2*^*−/−*^ and CCD treated control oocytes (Extended Data Table 2). First, this suggested that the physical properties of the nuclear envelope are similar in all cases. Second, this analysis showed that the power spectra of *Fmn2*^*−/−*^ and CCD treated oocytes display significantly lower effective temperatures *T*_*e*_ (5 fold, see Fig. 5C, 5D, Extended Data Fig. 4B and 4C and Extended Data Table 2). Overall this strongly suggests that the nuclear envelope fluctuations are generated by Formin 2 induced cytoplasmic activity and are not due to inherent changes of nuclear envelope properties in the different types of oocytes.

### Actin promotes chromosome motion inside the nucleus

We reasoned that the activity generated by the F-actin mesh at steady state could potentially be transmitted to the chromatin inside the nucleus, as was observed for microtubule-induced nuclear envelope fluctuations of *Drosophila* embryos^33^. On physical grounds, the active fluctuations of the nuclear envelope, induced by cytoplasm activity, exert fluctuating forces on the nuclear fluid and therefore on the chromatin. It is therefore expected that the chromatin is subjected to an active temperature, induced by cytoplasmic activity and transduced by the nuclear envelope. Assuming that the nucleoplasm behaves as a viscous fluid, a simple physical model (see Supplementary model) then predicts that the diffusion coefficient of a given tracer particle in the nucleus can be written *D* = *k*_*B*_(*T* + *αT*_*a*_)/*λ*, where *λ* is a friction coefficient, *T*_*a*_ models the active fluctuations of the cytoplasmic F-actin network introduced above, and *α* < 1 is a numerical constant that accounts for the dampening of these active forces in the nucleus. In this picture, the chromatin experiences an effective temperature *T*_*e (nucleoplasm)*_ = *T* + *αT*_*a*_, which depends on the cytoplasmic activity.

To test this model, we followed chromatin motion at high temporal resolution. A time projection of 10 time-points from the movie shows that in controls, chromatin is more mobile than in *Fmn2*^*−/−*^ oocytes (Fig. 6A compare upper and lower time projections and Supplementary Videos 7 and 8). The tracks of the nucleolus centroid show that the nucleolus explores more space in nuclei from controls than from *Fmn2*^*−/−*^ oocytes (Fig. 6B compare green to red tracks), in agreement with a higher effective temperature in controls. This was confirmed by the curve of the Mean Square Displacement (MSD), a measure of the space explored by an object per unit of time, of the nucleolus centroid which was always higher in controls than in *Fmn2*^*−/−*^ oocytes (Fig. 6C compare green with red curves). The MSD plots increased linearly with time, typical of a diffusive motion^34^, which argued that the nucleoplasm can be described as a viscous fluid. This is consistent with our proposed model. It also suggests that the intrinsic properties of nuclei are comparable between all types of oocytes. We addressed this further by performing intra-nuclear FRAP experiments of our nucleo-plasmic probe^13^, which is not associated with and thus diffuses independently of chromatin^35^. Fluorescence recovery was identical in *Fmn2*^*+/−*^ and *Fmn2*^*−/−*^ nuclei, supporting the finding of similar intrinsic nucleo-plasmic properties between the two conditions (Fig. 7A and B). The diffusion coefficient of the nucleolus, D, is proportional to the MSD (D = MSD/2dt, where d is the dimension of the chromatin and t is the time). Thus, the fit of the MSD curves showed that the diffusion coefficient of the nucleolus depended on the cytoplasmic F-actin since it was 3 times larger in *Fmn2*^*+/−*^ than in *Fmn2*^*−/−*^ or than in *Fmn2*^*+/−*^ oocytes treated with CCD for 1 h 30 (Extended Data Fig. 6A, B, and C and Supplementary Video 9). This observation is consistent with our physical model and suggests an effective temperature in the nucleoplasm (*T*_*e (nucleoplasm)*_), 3 times larger in controls. It is noteworthy that the effective temperature *T*_*e (nucleoplasm)*_ experienced by the chromatin is smaller than the effective temperature experienced by the nuclear envelope *T*_*e (cytoplasm)*_. This can be attributed to the dampening of active forces, generated at the nuclear periphery by the cytoplasmic F-actin network, within the nucleus.

**Figure 6:**
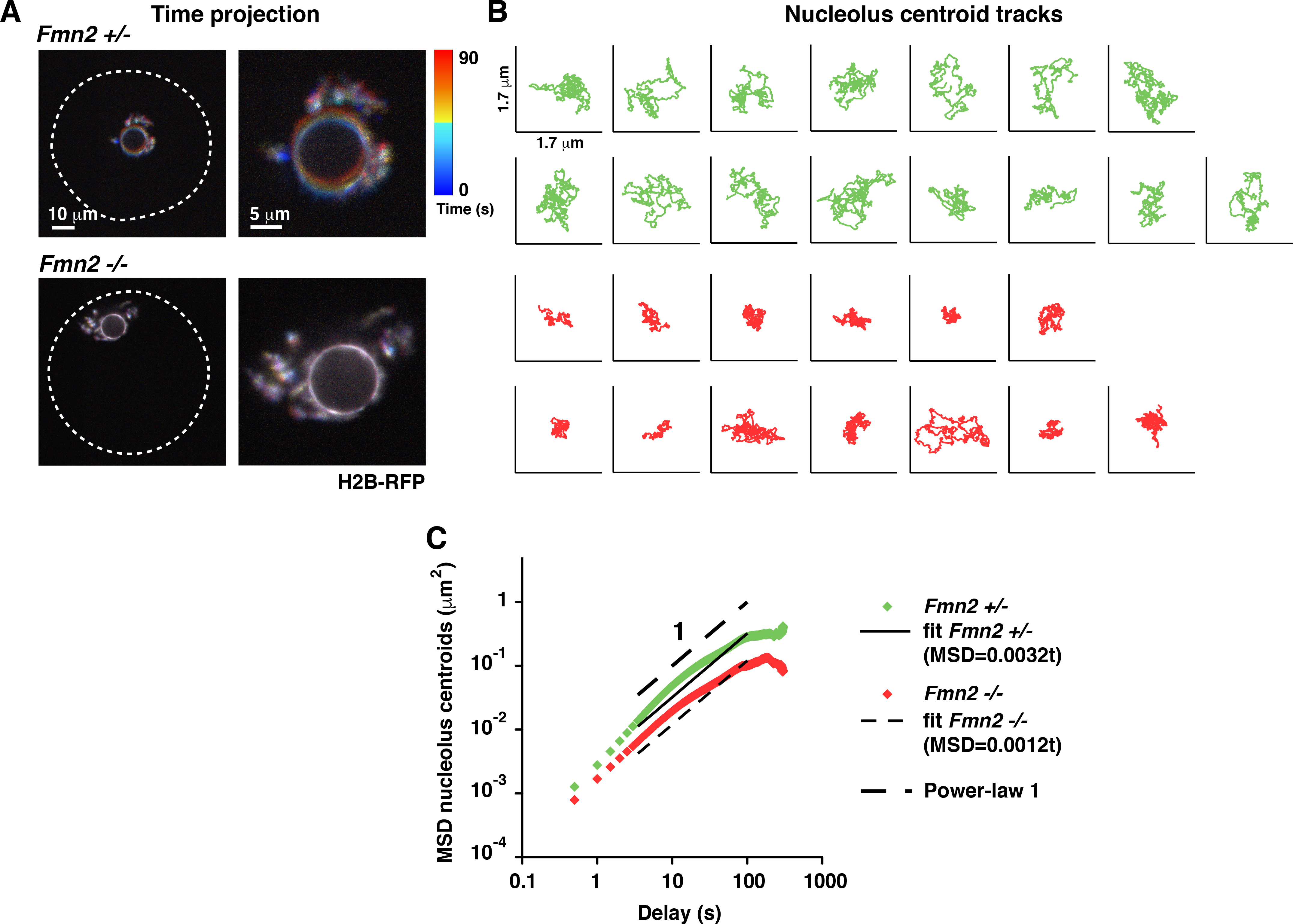
Nuclear envelope fluctuations mediate chromatin motion. **A:** Chromatin motion in oocytes with higher (*Fmn2*^*+/−*^) or lower (*Fmn2*^*−/−*^) nuclear envelope fluctuations. Top left: One movie coloured time-projection on 10 time-points (90 seconds) of a *Fmn2*^*+/−*^ oocyte injected with cRNAs coding for the chromatin probe H2B-RFP. Blue first timepoint, red last timepoint. The white dotted line represents the outline of the oocyte. Scale bar is 10 μm. Top right: Zoom of the chromatin area from the oocyte presented on the left panel. Scale bar is 5 μm. Bottom left: One movie coloured time-projection of a *Fmn2*^*−/−*^ oocyte injected with cRNAs coding for the chromatin probe H2B-RFP. Blue first timepoint, red last timepoint. The white dotted line represents the outline of the oocyte. Scale bar is 10 μm. Bottom right: Zoom of the chromatin area from the oocyte on the left. Scale bar is 5 μm. **B:** The nucleolus explores more space in *Fmn2*^*+/−*^ versus *Fmn2*^*−/−*^ oocytes. Top (green): Individual centroid tracks of 15 distinct nucleoli from *Fmn2*^*+/−*^ oocytes. Bottom (red): Individual centroid tracks of 13 distinct nucleoli from *Fmn2*^*−/−*^ oocytes. Total duration of the tracking is 5 minutes. **C:** Oocytes with higher nuclear envelope fluctuations display more important chromatin motion. Nucleoli from *Fmn2*^*+/−*^ and *Fmn2*^*−/−*^ display diffusive-like motion. Higher MSD levels in *Fmn2*^*+/−*^ reveal increased intra-nuclear activity. Mean MSD plot of nucleoli centroids in *Fmn2*^*+/−*^ (green) and *Fmn2*^*−/−*^ oocytes (red) presented as a function of the delay (in s) on a log/log scale. Approximate power-law slope of 1 is indicated. The shift between *Fmn2*^*+/−*^ and *Fmn2*^*−/−*^ nucleoli plots corresponds to a MSD ratio of 3. MSDs data are fitted to a straight line, yielding up MSD = 0.0032 *t* for *Fmn2*^*+/−*^ (plain line) and MSD= 0.0012 *t* for *Fmn2*^*−/−*^ (dotted line). *Fmn2*^*+/−*^: N=15 nucleoli from one experiment; *Fmn2*^*−/−*^: N=13 nucleoli from one experiment.

**Figure 7:**
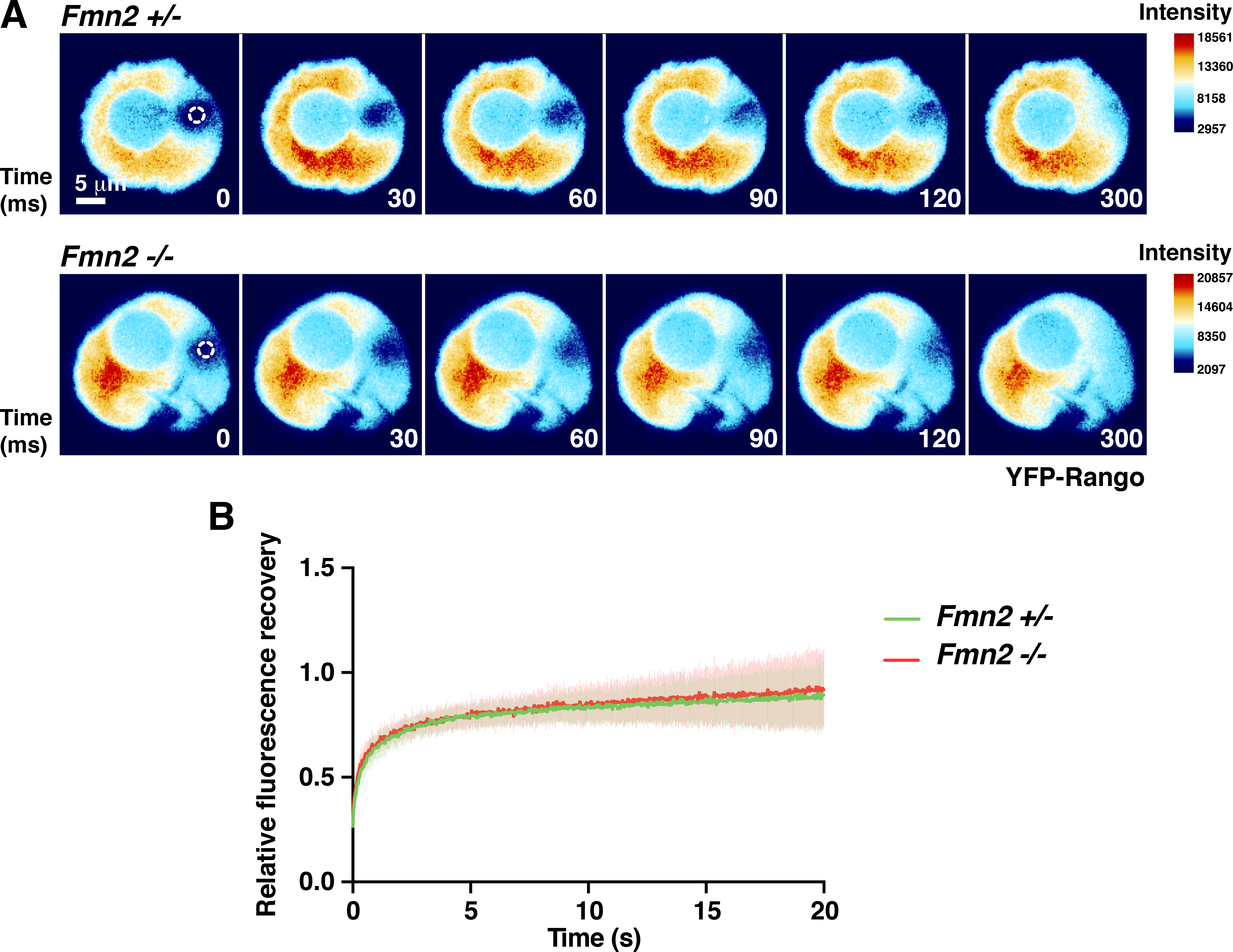
Identical biophysical nucleo-plasmic properties of *Fmn2*^*+/−*^ and *Fmn2*^*−/−*^ nuclei revealed by FRAP. **A:** Representative sequence of a FRAP experiment of the nuclear probe YFP-Rango in *Fmn2*^*+/−*^ (top) and *Fmn2*^*−/−*^ (bottom) nuclei. The white dotted circle represents the region where the signal recovery is measured (region of lowest total intensity at the beginning of the sequence, spot size diameter 2 μm). The color codes of the heat-maps represent the intensity of the signal. Blue: lower intensity level. Red: higher intensity level. Scale bar is 5 μm. **B:** Mean recovery curves of the YFP-Rango signal in *Fmn2*^*+/−*^ and *Fmn2*^*−/−*^ nuclei. Normalized fluorescence recovery corrected for bleaching (see Methods for a more detailed explanation) is plotted as a function of time for *Fmn2*^*+/−*^ (green) and *Fmn2*^*−/−*^ (red) nuclei. Error bars represent SD. N=23 oocytes for *Fmn2*^*+/−*^, 2 independent experiments and N=26 oocytes for *Fmn2*^*−/−*^, 3 independent experiments.

### Nuclear envelope and chromatin movements are correlated

Our model was further supported by the measure of the correlation between nuclear envelope and chromatin movements. To quantify this correlation over time, we measured the distance X from the nucleus centre of mass to the nuclear envelope outline and Y from nucleus centre of mass to the chromatin outline. Then the mean distance and the correlation between X and Y were calculated over time in all directions in the focal plane (i.e.: for all points of the nuclear circumference scanned using a revolving angle θ of 2° increment from 0 to 360°; Fig. 8A and see Methods). We established a correlation between nuclear envelope movements and chromatin movements in both *Fmn2*^*+/−*^ and *Fmn2*^*−/−*^ nuclei (Fig. 8B). The mean correlation was similar in both cases, again arguing that intra-nuclear properties are comparable in both types of oocytes (Fig. 8B). We noticed that the strength of this correlation remarkably increased when nuclear envelope and chromatin were close to one another (Fig. 8C, right panels, Supplementary Videos 10 and 11), in agreement with the model that hypothesizes that nuclear envelope acts as a source of fluctuations for the chromatin. The strength of this correlation is particularly high, as assessed by the values of correlation levels close to 1 obtained for the pooled *Fmn2*^*+/−*^ and *Fmn2*^*−/−*^ dataset when chromatin and nuclear envelope are closest to one other (Extended Data Fig. 7). This is a good illustration of the first law of geography by Waldo Tobler^36^, stating that “everything is related to everything else, but near things are more related than distant things.”

**Figure 8:**
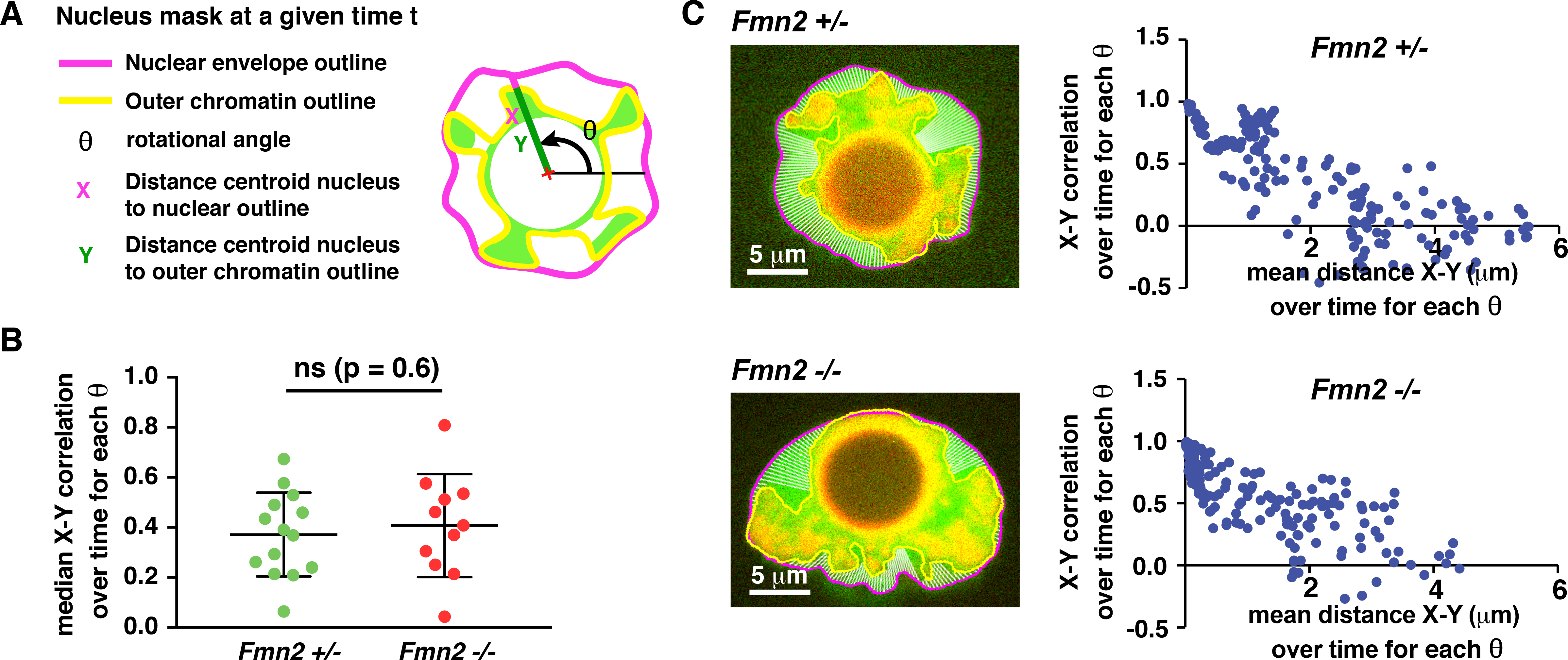
Correlation between nuclear envelope and chromatin motion in *Fmn2*^*+/−*^ and *Fmn2*^*−/−*^ nuclei. **A:** Scheme of the method used to quantify the correlation between nuclear envelope and chromatin dynamics over time and in a given direction. The nucleus is labelled with the nuclear probe YFP-Rango and the chromatin with H2B-RFP. Directions were defined by a revolving angle θ of 2° increment from 0 to 360°. For each direction, the distances X from the nucleus centroid to the nuclear envelope outline and Y from the nucleus centroid to the chromatin outline were computed. Then the mean distance and the correlation between X and Y were calculated over time for each θ. **B:** Nuclear envelope dynamics correlates similarly with chromatin dynamics in *Fmn2*^*+/−*^ and *Fmn2*^*−/−*^ nuclei. The median correlation over time of all θ is plotted for each nucleus. Error bars represent SD. Mean and SD are 0.37 +/− 0.17 μm for *Fmn2*^*+/−*^ and 0.41 +/− 0.21 μm for *Fmn2*^*−/−*^. P-values for Student t test is 0.6. N=14 oocytes for *Fmn2*^*+/−*^, one experiment and N=11 oocytes for *Fmn2*^*−/−*^, one experiment. **C:** Correlation between nuclear envelope and chromatin motion increases when chromatin and nuclear envelope are close. Top left: Example of *Fmn2*^*+/−*^ nucleus segmentation for further analysis with the method described in A. Top right: Correlation between X and Y over time for each θ as a function of the mean distance between X and Y over time for each θ for the nucleus presented in the left panel. 180 revolving angles every 2°. Bottom left: Example of *Fmn2*^*−/−*^ nucleus segmentation for further analysis with the method described in D. Bottom right: Correlation between X and Y over time for each θ as a function of the mean distance between X and Y over time for each θ for the nucleus presented in the left panel. 180 revolving angles every 2°. Scale bars are 5 μm.

To summarize, our work supports a mechano-transduction model where F-actin dependent cytoplasmic activity generates non-thermal fluctuations of the nuclear envelope, which in turn significantly enhances chromatin diffusion. This model is in agreement with gene expression data, where abolishing active fluctuations by a 5h-CCD treatment in control oocytes, which maintain central nuclei, leads to up-regulation of Cdc5L similarly to *Fmn2*^*−/−*^ oocytes (Extended data Fig. 3C).

## Discussion

In mouse oocytes, Formin 2 has been so far mainly investigated as a straight F-actin nucleator required for chromosome positioning^11–14,37–41^. We show that its role goes beyond and that Formin 2 is also involved in chromatin dynamics and regulation of gene expression. Formin 2 does not modulate the nuclear actin pool or the MAL-SRF activation pathway. We present evidence that Fmn2-assembled microfilaments in the cytoplasm exert fluctuating forces (due to an increase in effective temperature, 5 fold above thermal fluctuations) on the nucleus that are transmitted to the chromatin. When cumulated, these forces will eventually impact both chromatin motion as well as nuclear architecture. Much has been learned on how forces from the cell surroundings modulate the cell internal geometry, in particular how cytoskeletal elements transduce these external forces to organelles or to the mitotic spindle^42–46^. Adherent cells respond to mechanical forces of their environment and artificial forces applied to integrins are transduced to the nucleus, impacting its morphology^47^. The up-regulation of transcription of a multi-copy insertion of a Bacterial Artificial Chromosome (BAC) into cells in culture can be induced by integrin mediated local shear stress using magnetic twisting cytometry^48^.The process by which external forces are transduced to the nucleus is called mechano-transduction and its impact is summarized in this review^49^. We are presenting an original model of mechano-transduction, uncoupled from external forces, that could be extended to other isolated models, like early embryos, where the cell produces its own internal forces affecting nuclear position, shape, state of chromatin condensation and gene expression. Our rescue experiments show, in a physiological context, that up-regulation of endogenous gene transcription can be triggered via actin-based nucleus re-positioning in mouse oocytes. We demonstrate, with α-amanitin treatment, that this phenomenon goes through the modulation of the oocyte endogenous transcriptional response. Since we can detect a signature of genes up and down-regulated in *Fmn2*^*−/−*^ oocytes, we can imagine that some genes respond to mechano-transduction. Based on the result of the CCD treatment of control oocytes, which keep a central nucleus, we would like to argue that it is more the mechanism of positioning than the nucleus position itself which is important here. It will be interesting in future work to analyse in greater details those genes that respond to mechano-transduction and their relation with Lamin A ^15,16^.

In all organisms, the stockpile of maternal transcripts controls the capacity of the female gamete to turn into a viable embryo. Indeed and as we show here, all oocytes undergo a massive transcriptional silencing and it is only after fertilization, at various stages depending on the species, that zygotic transcription will be turned on^50^. Recent studies provided evidence that the quality of the oocyte maternal transcriptome controls the capacity to switch on zygotic genome activation after fertilization^51–53^. Therefore we propose that the forces that permit mouse oocyte nucleus centring modulate gene expression via a mechano-transductive effect and that this in turn is essential for the female gamete ability to transform into a developmentally competent embryo after fertilization. Importantly, an off-centre nucleus correlates with poor outcome for mouse and human oocyte development^7,8^. Oocytes are routinely manipulated for ART (Assisted Reproduction Technics), specifically in Western Countries where women tend to postpone childbearing. Furthermore, women undergoing chemotherapy treatments or women sensitive to hormonal treatments used for superovulation can depend on *in vitro* maturation of oocytes to conceive children. These ART techniques are still associated with significant risks. The criteria of central nucleus, visible by transmitted light, provides a non-invasive way to select for the most promising oocytes. Also, we have previously shown that a similar mechanism is at play in the one-cell-stage zygote to promote pronuclei centring^54^. Monitoring the extent of cytoplasmic flux, indirectly induced by the motion of actin-positive vesicles either in oocytes (for MIV) or in one-cell zygote (in FIV or ICSI) are rapid non-invasive (done by transmitted light) techniques that could provide a quantitative measure of oocyte and embryo fitness (as was done in^55^ and as we did both in^11,54^). Combining various non-invasive quantitative methods such as nucleus position and extent of cytoplasmic flux could benefit the selection of oocytes and embryos.

## Methods

### Oocyte collection, culture and microinjection

Oocytes were collected from 11 week-old OF1 and 15 weeks-old *Fmn2*^*+/−*^ or *Fmn2*^*−/−*^ female mice as previously described^56^ and maintained in Prophase I arrest in M2+BSA medium supplemented with 1 μM Milrinone^57^. At this stage, they were microinjected with cRNAs using an Eppendorf Femtojet microinjector. Oocytes were kept a minimum amount of time in Prophase I to allow sufficient expression of the various probes. It was respectively between 1 and 2 hours for YFP-Rango or H2B-RFP, 4 hours for RPEL1-eGFP-NLS3 and 2h30 for FRAP experiments on YFP-Rango. The expression time was identical for all types of oocytes in the various settings. For rescue experiments, *Fmn2*^*−/−*^ oocytes were kept in Prophase I for 8 hours after microinjection of Fmn2, FH1-FH2 or Nter cRNAs. This duration ensured that most oocytes completed the oocyte centring process (which is about 5 hours^11^). For rescue experiments with Nter cRNAs, where the nucleus does not recentre, oocytes were checked for Nter-eGFP fluorescence as an injection control. For detection of nascent transcripts, Prophase I arrested oocytes were incubated for 4 hours with 100 μM of the nucleoside analog 5’-ethynyl uridine (EU) from the Click-IT RNA Alexa Fluor 488 Imaging kit (Thermofisher, Ref. C10329). To block transcription, oocytes were pre-incubated for 1 hour in M2+BSA medium supplemented with 1 μM Milrinone +/− 75 μg ml^−1^ α-amanitin before proceeding to other treatments (microinjection for rescue experiments). All live culture and imaging were carried out under oil at 37°C. The animal facility scientific council of the CIRB granted permission for the animal experiments described here.

### Plasmids and *in vitro* transcription of cRNAs

We used the following constructs: pCS2-Fmn2-Myc and pCS2-Fmn2-eGFP (gifts from Philip Leder with modifications on amino acids sequence S944→P, L946→P, V1040→M, L1142→P according to Formin 2 protein sequence NP_062318.2), pRN3-YFP-Rango-CFP^35^, pRN3-H2B-RFP^58^. pCS2-Nter-eGFP was constructed by cloning the N-terminal domain of Formin 2 (amino acids 1 to 734) amplified from the previously described pCS2-Fmn2-Myc, into pCS2-eGFP. pCS2-FH1-FH2-eGFP was constructed by cloning the N-myristoyl domain MGNQDGK of Formin 2 (amino acids 1 to 7) and linker GGSGGGSG connected to the FH1-FH2 domain (amino acids 734 to 1578), amplified from the previously described pCS2-Fmn2-eGFP, into pCS2-eGFP. pRN3-RPEL1-eGFP-NLS3 was constructed by cloning the RPEL1-eGFP-NLS3 cassette into pRN3, amplified from RPEL1-eGFP-NLS3 in a pEGFP-N1 backbone (gift from Dyche Mullins, ^29^). The mcherry-Plk4 construct (gift from Renata Basto, ^59^) was subcloned into pRN3. *In vitro* synthesis of capped cRNAs was performed as previously described^60^. cRNAs were centrifuged at 4°C during 45 min at 13000 rpm before microinjection.

### Drug treatments

Cytochalasin D (Life Technologies, Ref. PHZ1063) was diluted at 10 mg ml^−1^ in DMSO and stored at −20°C. It was used on oocytes at 1 μg ml^−1^. Nocodazole (Sigma, Ref. M1404) was diluted at 10 mM in DMSO and stored at −20°C. It was used on oocytes at 1 μM. α-amanitin (Sigma, Ref. A2263) was diluted at 1 mg ml^−1^ in RNAse free water. It was used on oocytes at 75 μg ml^−1^.

### Oocyte RNA extraction, cDNA synthesis and quantitative PCR

Oocytes were collected as described above. Total RNA was extracted from oocytes (freshly collected or cultured; approximately 25 oocytes sample) using the RNAqueous®-Micro Total RNA Isolation Kit (Thermofisher, Ref. AM1931) following the manufacturer’s instructions. RNA was eluted in 20 μM of elution buffer. SuperScript II kit (Thermofisher, Ref. 18064014) was used for reverse transcription following the manufacturer’s instructions and using random primers. The cDNA was then used for quantitative PCR using LightCycler 480 (Roche) using the primer pairs listed (Supplementary Notes, Table 3; Actin b as a loading control). Relative expression levels were calculated using the 2-ΔΔC.

For confirmation of the transcriptomic analysis of *Fmn2*^*+/−*^ compared to *Fmn2*^*−/−*^ (Fig. 2C), mRNA quantity is normalized to Beta-actin (Actb). Mean and SEM are both normalized relative to *Fmn2*^*+/−*^. For rescue experiments (Fig. 3E and Extended Data Fig. 2), the expression relative to Beta-actin is calculated for non-injected and injected oocytes in each experiment. Then the mean and SEM of all experiments are calculated and the mean and SEM of both groups are divided by the mean of the non-injected oocytes (mean of the non-injected oocytes = 1).

### RNA extraction for RNAseq

Oocytes were collected as described above. Competent oocytes were selected following morphological criteria (size, zona thickness, perivitelline space). Total RNA was extracted from oocytes (freshly collected; 2 samples of *Fmn2*^*+/−*^, 50 oocytes each and 2 samples of *Fmn2*^*−/−*^, 50 oocytes each) using the RNAqueous®-Micro Total RNA Isolation Kit (Thermofisher, Ref. AM1931). Oocytes were washed 3 times in PBS, resuspended in lysis buffer, freezed in liquid nitrogen and conserved at −80°C overnight. After unfreezing, RNA extraction was carried out following the manufacturer’s instructions. RNA was eluted twice in 10 μl elution buffer. Samples were then treated with DNAse I.

### cDNA libraries and RNA sequencing

1 ng of total RNA were amplified and converted to cDNA using NuGEN’s Ovation RNA-Seq kit V2. Following amplification, 1 μg of cDNA was fragmented to approximately 200 bps using Covaris S200. The remainder of the library preparation was done using 200 ng of cDNA following TruSeq RNA Sample Prep v2 kit from the End Repair step. Libraries were multiplexed by 4 on 1 flow cell lane. A 50 bp read sequencing was performed on a HiSeq 1500 device (Illumina). A mean of 17.3 ± 3.9 million passing Illumina quality filter reads was obtained for each of the 4 samples. For each biological sample, 3 technical replicates were done.

### Bioinformatics analysis

Raw reads passing Illumina quality filters were cleaned of adapters, and mapped with STAR^61^. For each sample, between 4 and 15 millions reads were mapped at a unique location on the *Mus musculus* genome (mm10 GRCm38, with Ensembl83 annotation). Read counts per gene were also generated by STAR, and the DESeq2 package^62^ was used for differential expression analysis. Differentially expressed genes with an adjusted p-value threshold of 0.05 were selected for further analysis.

### Gene Ontology analysis

Gene ontology enrichment analysis for Biological Processes was performed on the Gene Ontology Consortium website against *Mus musculus* reference list using the PANTHER over-representation test with Bonferroni correction.

### Live imaging

Spinning disk images were acquired at 37°C in M2 + BSA +1 μM Milrinone using a Plan-APO 40x/1.25 NA objective on a Leica DMI6000B microscope enclosed in a thermostatic chamber (Life Imaging Service) equipped with a Retiga 3 CCD camera (QImaging) coupled to a Sutter filter wheel (Roper Scientific) and a Yokogawa CSU-X1-M1 spinning disk. Metamorph software (Universal Imaging, version 7.7.9.0) was used to collect data.

For imaging the nucleus labelled with YFP-Rango, oocytes were illuminated with an excitation wavelength of 491 nm every 500 ms with the stream acquisition mode of Metamorph on one single plane focused on the nucleolus. For imaging chromatin labelled with H2B-RFP, oocytes were illuminated with an excitation wavelength of 561 nm every 500 ms with the stream acquisition mode of Metamorph on one single plane focused on the nucleolus. For RPEL1-eGFP-NLS3 imaging together with H2B-RFP, images were acquired with a respective excitation wavelength of 491 nm and 561 nm during 500 ms. Z-series were performed with Z-steps of 1 μm. To study the correlation between nuclear envelope movements and chromatin movements, YFP-Rango and H2B-RFP were imaged together in the following conditions: images were acquired every 30 s with a respective excitation wavelength of 491 nm and 561 nm during 500 ms. In these conditions, the time shift between the two channels is 850 ms, only 2,8% of the 30 s time interval between two successive time points. For mcherry-Plk4 imaging together with YFP-Rango, images were acquired with a respective excitation wavelength of 561 nm and 491 nm during 500 ms. Z-series were performed with Z-steps of 0.5 μm.

The FRAP experiments were done at 37 °C in M2 + BSA +1 μM Milrinone on a Nikon Eclipse TL microscope enclosed in a thermostatic chamber equipped with an Evolve EMCCD camera (Photometrics) coupled to a Sutter filter wheel (Roper Scientific) and a Yokogawa CSUX1-A1 spinning disc. FRAP experiments were performed on oocytes expressing the YFP-Rango nuclear probe. For the FRAP routine, an Ilas2 targeted laser illumination system (Roper Scientific) was used. 491 nm full power laser line was activated during 202 ms in bleach point mode with a Plan Apo lambda 60X, N.A: 1.4 (Nikon). 6 mW power at 491 nm was measured at back focal plane. The recovery sequence was realized by acquiring 256×256 pixels images, with 33 ms exposure time at maximum frame rates (30 frames per second), using 10% laser power.

### Immunofluorescence

Oocytes were fixed 30 min at 30°C in 4% paraformaldehyde on coverslips treated with gelatin and polylysine. Permeabilization was done by incubating oocytes in 0.5% Triton-X in PBS for 10 min at room temperature. For Lamin A staining, a rabbit monoclonal antibody was used against Lamin A (Abcam, clone (EP4520-16), Ref. ab133256) at 1:500 dilution. A Cy3-conjugated secondary antibody (Jackson Immuno Research, Ref. 711-165-152) was used at 1:150 dilution. EU detection was carried out following the instructions of the Click-IT RNA Alexa Fluor 488 Imaging kit (Thermofisher, Ref. C10329). DNA was labelled with DAPI in Prolon Gold mounting medium (Thermofisher, Ref. P36941). Oocytes were mounted onto 250 nm thick perforated stickers to avoid smashing of the oocytes (Electron Microscopy Sciences, Ref. 70366-12).

For Lamin A and DAPI staining, images were acquired using a Leica SP5 confocal microscope with a Plan APO63/1.25NA objective. Z-series were performed with Z-steps of 0.5 μm. For EU signal imaging, images were acquired using a Leica DMI6000B microscope equipped with a Retiga 3 CCD camera (QImaging) coupled to a Sutter filter wheel (Roper Scientific) and a Yokogawa CSU-X1-M1 spinning disk, with a Plan-APO 40x/1.25 NA objective.

### Image analysis

When the shape of nuclei was asymmetric (for *Fmn2*^*−/−*^ and *Fmn2*^*+/−*^ + CCD), nuclei labelled with YFP-Rango were rotated on Fiji^63^ to orient the smooth part up and the invagination down, in order to analyse nuclear envelope fluctuations. Then a custom plugin developed for Fiji was used to remove the noise in the signal using the Pure denoise plugin, threshold the signal and fill the hole corresponding to the nucleolus in order to create a binary nucleus mask, realign it and calculate the distance r from the centroid of the nucleus mask to the circumference of the mask for all θ angles (θ from 0 to 360° by 1° increment, as defined in Fig. 5B). Results were provided as xls files. Then, for each nucleus, the mean distance R over all the 600 time points t for each defined θ angle was calculated, allowing to plot the mean shape over time (Fig. 5B). For each θ, the mean distance R was subtracted from the distance r for all time points t. The variance (r-R)^2^ is a measure of nuclear envelope fluctuations. The mean fluctuation of nuclear shape was calculated for all time points t and all angles θ. Eventually, for all nuclei coming from one condition (*Fmn2*^*+/−*^, *Fmn2*^*−/−*^ and *Fmn2*^*+/−*^ + CCD), the mean of the fluctuations for all time points t and all angles θ was calculated.

Heatmaps representing the mean fluctuations for each defined θ angle over all time points t and plotted along the contour of the mean shape were generated using the G-plot package on R (version 3.3.2).

Fourier transform calculation was done on the fluctuations values (r-R) for all angles θ and all time points t. We used the fft package on R to extract the values of the Fourier transform moduli.

Nucleolus automated tracking was performed on the same movies coming from nuclei labelled with YFP-Rango used to analyse nuclear envelope fluctuations, similarly reoriented with the smooth part up and the invagination down. A custom plugin for Fiji was developed to threshold the signal and fill the hole corresponding to the nucleolus in order to create the binary nucleus mask and realign it. The mask of the nucleolus was then extracted from the realigned nucleus mask, allowing the tracking of the centroid of the nucleolus in the frame of the realigned nucleus. Trajectories were provided as xls files and then converted to xml files for Mean Squared Displacement analysis.

For time projections of chromatin movies labelled with H2B-RFP, the background was subtracted and bleaching was corrected using the Histogram Matching algorithm on Fiji. A sub-stack of 10 planes every 10 s (90 s total duration) was selected and time-projection was done with the Temporal-colour code plugin on Fiji, using a custom LUT.

For Mean Square Displacement analysis of nucleoli trajectories, we used the@msdanalyzer class of MATLAB (version 2016a) described in: http://bradleymonk.com/matlab/msd/MSDTuto.html

Single plane images of nuclei stained for Lamin A and DAPI (Fig. 1A and 3B) were done on Fiji.

EU signal analysis was done using Metamorph (Extended Data Fig. 3B). After substracting the backround, the integrated intensities of EU and DAPI staining were determined in the mid-sectional plane, within the same ROI for EU and DAPI channels for a given oocyte. ROIs of identical areas were used for all oocytes in all conditions. The EU:DAPI ratio was calculated.

RPEL1 signal analysis (Fig. 4C) was done on Metamorph. Planes below and above the nucleus were removed from Z-stacks. A sum projection was done on the remaining stacks and the background was subtracted from the resulting image. The total intensity was calculated in a region including the nucleus. The same region was used for RPEL1-eGFP-NLS3 and for H2B-RFP. The same region area was used for all nuclei.

To determine the nucleo-cytoplasmic ratio of YFP-Rango (Extended Data Fig. 1), after background subtraction, total intensities were determined on Metamorph in 6 circular regions of the same size (2 in the nucleus and 4 in the cytoplasm). The mean of nuclear intensities and cytoplasmic intensities and the corresponding ratio were calculated for each nucleus.

For FRAP analysis, after background subtraction, a circular region of 2 μm diameter, the same as the spot size, was positioned in the region of lowest total intensity in the first frame of the post-bleach sequence. Total intensity in this region was then measured on Metamorph for all the time points of the sequence. In order to be able to compare different experiments, the last pre-bleach time point was normalized to 1. The normalized data curves were corrected for the photo-bleaching occurring during acquisition by dividing the normalized fluorescence by a function modeling photo-bleaching. This function was determined by fitting the fluorescence decay observed in the pre-bleach phase to a mono-exponential function of the type F(t) = (Fi- Finf) e^−t/ τ^ + Finf, where Fi is the normalized fluorescence intensity at the first pre-bleach time point, Finf is the normalized fluorescence intensity at infinite and τ is the time constant corresponding to the time when 63% of the total photo-bleaching had occurred. Fi, Finf and τ were determined by the fitting.

### Computational 3D imaging

<u>3D image segmentation</u>: in order to use of isometric 3D kernels, we performed a straightforward anisotropy correction of the stack of images by performing a cubic interpolation in the z direction. Thus, the interpolated stack of images have equal dimensions on x, y and z axes. After denoising using a Laplacian filtering (sigma=3), a robust 3D segmentation of the two channels of the 3D stack image was performed using a k-means clustering of the grey levels and spatial information^64^ implemented in the scikit-image python package (version 0.11.3 of Python 2.7). Each channel is split into three clusters. The cluster with the highest intensity corresponds to the foreground object in each channel (Lamin A or DNA). The segmentation algorithm considers a “compactness” parameter that balances color. Spatial relationships was set to 0.01 to segment the DNA signal and 0.005 to segment the Lamin A signal properly. The parameter was then maintained unchanged for the whole dataset.

<u>3D features extraction</u>: 94 quantitative features were extracted from DNA and Lamin A foregrounds to describe the morphological changes for all stacks in all conditions. The full list of features is shown in Supplementary Methods. Features can be summarized into shape and volume, DNA dispersion, features measuring the overlap between DNA and Lamin A signals, features measuring the isolated and detached pieces of DNA and features measuring spreadness and distances between DNA and Lamin A signals.

<u>Data analysis</u>: after extracting features using a computing cluster, each 3D image for each condition was represented by a vector of 94 features. Those data were then analyzed using two approaches:

*Univariate data analysis:* chosen features were used to compare selected couples of conditions using univariate statistical tests: Mann Whitney U test (one tailed) with Bonferroni correction when needed.

*Multivariate data analysis:* Highly correlated features were identified as connected components of a complete weighted correlation graph thresholded at 0.8 and the feature with the maximum of connections on each connected component was selected. This feature selection step retained 46 features (see Supplementary Methods). Those data were then analyzed using two approaches. Linear Discriminant Analysis^65^ was then performed on subsets of conditions using the 46 remaining features: each nucleus, now profiled as a 46-dimensional features was projected onto a lower-dimensional space by simultaneously maximizing the inter class variance and minimizing the intra class variance. The linear discriminant axes display the projection of the dataset that best separates the various oocyte conditions.

### Correlation of chromatin and nuclear envelope movements

This analysis was done using Python 2.7 using the scikit-image and scipy libraries. RFP (chromatin labelled with H2B-RFP) and GFP (nucleus labelled with YFP-Rango) image channels where smoothed with Gaussian kernel size 1.5 and 4 pixels respectively to suppress noise artefacts and signal was segmented using an automated Otsu threshold. Holes where filled and contours were extracted with a morphological gradient with a circular structural element of 3 pixels diameter. Aside, both channels were merged and smoothed with a Gaussian kernel of 90 pixels size and segmented with an automated Otsu threshold in order to compute a common centroid for each nucleus at each time point. Cartesian coordinates of each pixel of the contours of both channels were converted into polar coordinates with the centroid as origin for each image of each sequence. Contours were re-parametrised such that to obtain two radii values for each second degree direction: one for the chromatin contour X and one for the nucleus contour Y. At the end of the process, 180 values of X and 180 corresponding values of Y were obtained for each image of each video using the same set of parameters mentioned above. For each of the 180 angles, Spearman correlation coefficient between X and Y and mean distance between X and Y were computed over time. All images of all movies of both conditions were processed with the same parameter set.

### Non linear fitting of Fourier transforms

For each nucleus, squared values of Fourier transform moduli were averaged for each mode over all time points t. Nuclei whose shape was too far from a circular shape were excluded from the analysis (2 nuclei for *Fmn2*^*−/−*^, 2 for *Fmn2*^*+/−*^ + CCD and 4 for *Fmn2*^*−/−*^ + CCD). Data were fitted to the function 1/ (σ*n*^2^ + κ*n*^4^) + Yinf according to the Helfrich model. The additional off-set parameter Yinf was introduced to take into account the background high frequency noise in the data of the power spectrum. Data processing and fitting were done using the open source software R. For each nucleus, data were linearly interpolated using the function “approx”, for values of θ from 2° to 180°. Interpolated data were then averaged for each condition (*Fmn2*^*+/−*^, *Fmn2*^*−/−*^ and *Fmn2*^*+/−*^ + CCD, *Fmn2*^*+/−*^ + NZ, *Fmn2*^*−/−*^ + NZ). The model was finally fitted to the averaged data with the Marquardt method using the function “nlsLM” of the R package minpack.lm. Data for modes < 5 and > 178 were not considered for the fit because of lacking values except for *Fmn2*^*+/−*^. Squared values of Fourier transform moduli, averaged and predicted from the fitted model, where then exported as .csv files and plotted with Microsoft Excel.

### Code availability

For 3D computational analysis of nuclear architecture, all code needed to reproduce the results can be found at https://github.com/biocompibens/Meiospin.

For analysis of nuclear envelope fluctuations and tracking of the nucleoli, the code can be found at https://github.com/pmailly/OvocyteNucleusAnalyze.

### Data availability

The RNAseq data that support the findings of this study have been deposited in the Gene Expression Omnibus with the accession number GSE103718 (password apcfygoovtmrdwv).

### Statistical analysis

Mann-Whitney tests with Bonferroni correction were performed with Python 2.7 using the scipy.stats library (one-tailed tests) or with the GraphPad Prism software v 7.0 (two-tailed tests). Two-tailed student t tests were performed with Python or with Prism. One-way ANOVA followed by Tukey’s multiple comparison tests for ratios was performed with Prism. Linear fits of MSDs data were done using the Solver of Excel.

## Acknowledgments

We thank Cécile Sykes (Curie Institute), Adel Al Jord (CIRB) and Marie-Emilie Terret (CIRB) for critical reading of the manuscript. We thank all members of the Terret/Verlhac team for helpful discussions. Library preparation and Illumina sequencing were performed at the Ecole Normale Supérieure Genomic Platform (IBENS, Paris, France). We thank Dyche Mullins (UCSF) for the RPEL1-eGFP-NLS3 construct and Renata Basto (Curie Institute, Paris) for the mcherry-Plk4 construct. The work at IBENS Genomic Platform was supported by the France Génomique national infrastructure, funded as part of the “Investissements d’Avenir” program managed by the Agence Nationale de la Recherche (reference: ANR-10-INBS-09). This work was supported by grants from the Fondation pour la Recherche Médicale (FRM Label to MHV-DEQ20150331758), from the ANR (ANR-14-CE11 to MHV) and from the Labex Memolife (to MHV). This work has received support under the program « Investissements d’Avenir » launched by the French Government and implemented by the ANR, with the references: ANR-10-LABX-54 MEMO LIFE, ANR-11-IDEX-0001-02 PSL* Research University.

## Author Contributions

MA and MHV conceived and supervised the project. MA performed most experiments. MA and MHV analyzed most experiments. AO and AG conceived and did the 3D computational imaging approach. AG did the measure of correlation between nuclear outline and chromatin motion. TP assisted with FRAP experiments. SEH performed and analyzed the RT-qPCRs. For RNA seq, IQ did the RNA extraction, FC and SL did the cDNA libraries and performed the RNAseq, LB did the bioinformatics analysis. PM did the plugins to analyse nuclear envelope fluctuations and nucleolus tracking. CK did the fit for Fourier Transform analysis. RV designed the mathematical model. MA and MHV wrote the manuscript, which was seen and corrected by all authors.

## Author information

Reprints and permissions information is available at http://www.nature.com/reprints. The authors declare no competing financial interests. Correspondence and requests for materials should be addressed to.

## Extended Data Figure Legends

### Extended Data Figure 1: Nucleo-cytoplasmic ratio of the YFP-Rango nuclear probe suggests preservation of nuclear envelope integrity in *Fmn2*^*−/−*^ oocytes

Quantification of the nucleo-cytoplasmic ratio of YFP-Rango in *Fmn2*^*+/−*^ and *Fmn2*^*−/−*^ nuclei and *Fmn2*^*+/−*^ nuclei treated with 1 μg ml^−1^ CCD. Error bars represent SD. Mean and SD are 45.2 +/− 23.4 for *Fmn2*^*+/−*^, 57.9 +/− 23 for *Fmn2*^*−/−*^. Mann-Whitney U p-values is 0.063. N=15 for *Fmn2*^*+/−*^, one experiment, N=14 for *Fmn2*^*−/−*^, one experiment.

### Extended Data Figure 2: Rescue of gene expression levels by full-length Formin 2 is due to transcription, at least for the down-regulated genes

Rescue of gene expression defects of *Fmn2*^*−/−*^ oocytes by full-length Formin 2, addressed by RT-qPCR on 5 differentially expressed genes in *Fmn2*^*−/−*^, confirmed in 2C (4 downregulated and 2 upregulated), with or without treatment with 75 μg ml^−1^ of the transcriptional inhibitor α-amanitin. Mean of the normalized ratios of transcript levels between *Fmn2*^*−/−*^ and *Fmn2*^*−/−*^ + Fmn2 and between *Fmn2*^*−/−*^ and *Fmn2*^*−/−*^ + Fmn2 + α-amanitin. Error bars represent SEM. Mean of the ratios between *Fmn2*^*−/−*^ and *Fmn2*^*−/−*^ + Fmn2 and SEM are 0.85 +/− 0.090 (Acp1), 1.90 +/− 0.10 (Tcstv1), 1.33 +/− 0.046 (Tsc1), 0.80 +/− 0.19 (Cdc5L), 0.72 +/− 0.029 (H2eb1). Mean of the ratios between *Fmn2*^*−/−*^ and *Fmn2*^*−/−*^ + Fmn2 + α-amanitin and SEM are 0.82 +/− 0.10 (Acp1), 0.95 +/− 0.12 (Tcstv1), 0.89 +/− 0.055 (Tsc1), 0.96 +/− 0.086 (Cdc5L), 0.95 +/− 0.080 (H2eb1).

Statistics between *Fmn2*^*−/−*^ - *Fmn2*^*−/−*^ + Fmn2 (top of blue histogram), and *Fmn2*^*−/−*^ - *Fmn2*^*−/−*^ + Fmn2 + α-amanitin (black histogram) are presented. P-values for Tukey’s multiple comparison tests for ratios following one-way ANOVA are 0.4288 (Acp1), 0.0011 (Tcstv1), 0.0035 (Tsc1), 0.515 (Cdc5L) and 0.0155 (H2eb1) for *Fmn2*^*−/−*^ - *Fmn2*^*−/−*^ + Fmn2, 0.3234 (Acp1), 0.9376 (Tcstv1), 0.2301 (Tsc1), 0.9775 (Cdc5L) and 0.7332 (H2eb1) for *Fmn2*^*−/−*^ - *Fmn2*^*−/−*^ + Fmn2 + α-amanitin and 0.9674 (Acp1), 0.0008 (Tcstv1), 0.0008 (Tsc1), 0.6267 (Cdc5L) and 0.0379 (H2eb1) for *Fmn2*^*−/−*^ + Fmn2 - *Fmn2*^*−/−*^ - *Fmn2*^*−/−*^ + Fmn2 + α-amanitin. 3 independent experiments, N=20 oocytes by condition and by experiment.

### Extended Data Figure 3: Residual transcription in competent oocytes and kinetics of the impact of F-actin on Cdc5L expression in control oocytes

**A:** Residual transcription in fully grown (competent) oocytes. Transcriptionally active incompetent (left), competent (middle) and transcriptionally silent meiosis I (MI) oocytes (right) stained with DAPI (top, blue) with or without EU incorporation for 4 hours (bottom, red). Images were scaled differently to overcome differences in intensity ranges between conditions. One single plane is presented. Scale bar is 10 μm.

**B:** Quantification of the EU signal in competent compared to incompetent and to meiosis I oocytes. EU signal is normalized to DAPI. Error bars represent SD. Mean and SD for EU:DAPI ratio are 2.59 +/− 1.23 for incompetent + EU, 0.15 +/− 0.022 for incompetent - EU, 0.47 +/− 0.38 for competent + EU, 0.18 +/− 0.05 for competent - EU and 0.14 +/− 0.032 for MI + EU. N=14 for incompetent + EU, N=12 for incompetent - EU, N=17 for competent + EU, N=10 for competent - EU and N=14 for MI + EU. 2 independent experiments. Incompetent EU-incompetent no EU: Mann-Whitney U p-value = 3.50E-06. Incompetent EU-MI EU: Mann-Whitney U p-value = 1.49E-05. Competent EU-competent no EU: Mann-Whitney U p-value = 7.76E-03. Competent EU-MI EU: Mann-Whitney U p-value = 6.72E-05.

**C:** Determination of levels of a highly deregulated gene Cdc5L by RT-qPCR after cytoplasmic F-actin disruption by CCD. Mean of the normalized ratios of transcript levels between control oocytes and oocytes treated with 1 μg ml^−1^ CCD for 1, 3 and 5 hours. Error bars represent SEM. Mean and SEM are 0.92 +/− 0.066 (CCD 1h), 1.18 +/− 0.26 for (CCD 3h) and 3.99 +/− 0.54 (CCD 5h). P-values for t-test are: 0.308 (CCD 1h), 0.537 (CCD 3h) and 0.0052 (CCD 5h). 3 independent experiments, N=20 oocytes by condition and by experiment.

### Extended Data Figure 4: The cytoplasmic actin meshwork largely contributes to nuclear envelope fluctuations

**A:** Nuclear envelope fluctuations and nuclear shape are perturbed after treatment of control oocytes with CCD for 1 h 30. Left: images from a movie of a *Fmn2*^*+/−*^ oocyte injected with cRNAs coding for the nuclear probe YFP-Rango and treated with 1 μg ml^−1^ CCD. Right: Heat map of the mean nuclear outline over time (5 minutes) measured from the *Fmn2*^*+/−*^ oocyte treated with CCD presented on the left panel. The colour code represents the nuclear outline fluctuations over time (5 minutes), in μm^2^, for each position along the circumference relative to the mean shape over time. Red is for higher values of fluctuations and blue for lower values of fluctuations.

**B:** Nuclear envelope fluctuations are perturbed in both *Fmn2*^*−/−*^ and *Fmn2*^*+/−*^ + CCD oocytes. Nuclear fluctuations over 4 to 5 minutes, represented by the variance of the radius over t and θ, from *Fmn2*^*+/−*^, *Fmn2*^*−/−*^ and *Fmn2*^*+/−*^ + CCD nuclei. For *Fmn2*^*−/−*^ and *Fmn2*^*+/−*^ + CCD, the region of the large cleft was excluded from the measurements. Error bars represent SD. Mean and SD 0.34 +/− 0.13 μm^2^ for *Fmn2*^*+/−*^, 0.057 +/−0.038 μm^2^ for *Fmn2*^*−/−*^ and 0.07 +/− 0.04 for *Fmn2*^*+/−*^ + CCD. Mann-Whitney U p-values are < 0.0001 for *Fmn2*^*+/−*^ - *Fmn2*^*−/−*^, < 0.0001 for *Fmn2*^*+/−*^ - *Fmn2*^*+/−*^ + CCD and 0.2987 for *Fmn2*^*−/−*^ - *Fmn2*^*+/−*^ + CCD. N=33 oocytes for *Fmn2*^*+/−*^, 3 experiments, N=25 oocytes for *Fmn2*^*−/−*^, 3 experiments and N=14 for *Fmn2*^*+/−*^ + CCD, one experiment.

**C:** Power spectrum of nuclear fluctuations over 4 to 5 minutes: the squared moduli of the Fourier transforms of fluctuations (r-R) are plotted as a function of the mode for *Fmn2*^*+/−*^ (green), *Fmn2*^*−/−*^(red) and *Fmn2*^*+/−*^ + CCD (orange) nuclei. For *Fmn2*^*−/−*^ and *Fmn2*^*+/−*^ + CCD, the region of the large cleft was excluded from the measurements. Data are fitted to the function 1 / (σ*n*^2^ + κ*n*^4^) + Yinf according to the Helfrich model.

### Extended Data Figure 5: Microtubules reduce nuclear envelope fluctuations

**A:** The invagination in *Fmn2*^*−/−*^ nuclei contains a major aMTOC. Z-stacks of 1 μm of a live representative *Fmn2*^*−/−*^ nucleus. The aMTOC is labelled with mcherry-Plk4 (magenta) and the nucleus is labelled with the nucleo-plasmic probe YFP-Rango (green). Scale bar is 5 μm.

**B:** Treatment with NZ for 1h30 to 2 hours increases nuclear envelope fluctuations in control oocytes and affects nuclear shape in *Fmn2*^*−/−*^ oocytes. Left: images from movies of *Fmn2*^*+/−*^ and *Fmn2*^*−/−*^ oocytes injected with cRNAs coding for the nuclear probe YFP-Rango and treated or not with 1 μM NZ. Right: Heat map of the mean nuclear outline over time (4 minutes) measured from the oocytes presented on the left panel. The colour code represents the nuclear outline fluctuations over time (4 minutes), in μm^2^, for each position along the circumference relative to the mean shape over time. Red is for higher values of fluctuations and blue for lower values of fluctuations.

**C:** Nuclear envelope fluctuations are twice higher after NZ treatment in control oocytes. Nuclear envelope fluctuations over 4 to 5 minutes represented by the variance of the radius over t and θ from *Fmn2*^*+/−*^ and *Fmn2*^*−/−*^ nuclei. For *Fmn2*^*−/−*^, the region of the large cleft was excluded from the measurements. Error bars represent SD. Mean and SD are 0.34 +/− 0.13 μm^2^ for *Fmn2*^*+/−*^, 0.62 +/− 0.28 μm^2^ for *Fmn2*^*+/−*^ + NZ, 0.057 +/−0.038 μm^2^ for *Fmn2*^*−/−*^ and 0.073+/−0.064 μm^2^ for *Fmn2*^*−/−*^ +NZ. Mann-Whitney U p-values are 0.0002 for *Fmn2*^*+/−*^ - *Fmn2*^*+/−*^ +NZ and 0.89 for *Fmn2*^*−/−*^ - *Fmn2*^*−/−*^ +NZ. N=33 oocytes for *Fmn2*^*+/−*^, 3 experiments, N= 18 oocytes for *Fmn2*^*+/−*^ + NZ, 2 experiments, N=25 oocytes for *Fmn2*^*−/−*^, 3 experiments and N=15 for *Fmn2*^*−/−*^ + NZ, 2 experiments.

**D:** Power spectrum of nuclear fluctuations over 4 to 5 minutes: the squared moduli of the Fourier transforms of fluctuations (r-R) are plotted as a function of the mode for *Fmn2*^*+/−*^ (green), *Fmn2*^*+/−*^ +NZ (dark green), *Fmn2*^*−/−*^ (red) and *Fmn2*^*−/−*^ + NZ (brown) nuclei. For *Fmn2*^*−/−*^, the region of the large cleft was excluded from the measurements. Data are fitted to the function 1 / (σ*n*^2^ + κ*n*^4^) + Yinf according to the Helfrich model.

### Extended Data Figure 6: Treatment with CCD impairs chromatin motion

**A:** Nucleolar motion in *Fmn2*^*+/−*^ oocytes treated with CCD for 1 h 30. Left: One movie coloured time-projection on 10 time-points (90 seconds) of an *Fmn2*^*+/−*^ oocyte injected with cRNAs coding for the chromatin probe H2B-RFP and treated with 1 μg ml^−1^ CCD. Blue first timepoint, red last timepoint. The white dotted line represents the outline of the oocyte. Scale bar is 10 μm. Right: Zoom of the chromatin area from the oocyte on the left. Scale bar is 5 μm.

**B:** Individual centroid tracks of 17 nucleoli from *Fmn2*^*+/−*^ oocytes treated with CCD. Total duration of the tracking is 5 minutes.

**C:** Mean MSD plot of nucleoli centroids in *Fmn2*^*+/−*^ (green), *Fmn2*^*−/−*^(red) and *Fmn2*^*+/−*^ + CCD oocytes (orange). Approximate power-law slope of 1 is indicated. MSDs data are fitted to a straight line, yielding up MSD = 0.0032 *t* for *Fmn2*^*+/−*^ (plain line), MSD= 0.0012 *t* for *Fmn2*^*−/−*^ (dotted line) and MSD= 0.0014 *t* for *Fmn2*^*+/−*^ + CCD (small dotted line). *Fmn2*^*+/−*^: N=15 nucleoli in one experiment; *Fmn2*^*−/−*^: N=13 nucleoli from 13 oocytes in one experiment. *Fmn2*^*+/−*^ + CCD: N=17 nucleoli in one experiment.

### Extended Data Figure 7: Strong level of correlation between nuclear envelope and chromatin motion in *Fmn2*^*+/−*^ and *Fmn2*^*−/−*^ oocytes

Frequency distribution of the correlation values calculated following the method described in Figure 8A (see Fig. 8A and Methods). Correlations between X and Y over time were retrieved and plotted for all nuclei for the 20% of angles θ where the membrane was the closest to the chromatin. N=25 nuclei (N=14 oocytes for *Fmn2*^*+/−*^, one experiment and N=11 oocytes for *Fmn2*^*−/−*^, one experiment).

